# Balance between mutually exclusive traits shifted by variants of a yeast transcription factor

**DOI:** 10.1101/117911

**Authors:** Michael W. Dorrity, Josh T. Cuperus, Jolie A. Carlisle, Stanley Fields, Christine Queitsch

## Abstract

In *Saccharomyces cerevisiae*, the decision to mate or invade relies on environmental cues that converge on a shared transcription factor, Ste12. Specificity toward invasion occurs via Ste12 binding cooperatively with the co-factor Tec1. Here, we characterize the *in vitro* binding preferences of Ste12 to identify a defined spacing and orientation of dimeric sites, one that is common in pheromone-regulated genes. We find that single amino acid changes in the DNA-binding domain of Ste12 can shift the preference of yeast toward either mating or invasion. These mutations define two distinct regions of this domain, suggesting alternative modes of DNA binding for each trait. Some exceptional Ste12 mutants promote hyperinvasion in a Tec1-independent manner; these fail to bind cooperative sites with Tec1 and bind to unusual dimeric Ste12 sites that contain one highly degenerate half site. We propose a model for how activation of invasion genes could have evolved with Ste12 alone.

## Introduction

Transcription factors interact with DNA, with co-factors and with signaling proteins to allow cells to respond to changes in their environment. Despite the requirement to manage these multiple levels of regulation, most eukaryotic transcription factors possess a single DNA-binding domain. Distinct responses must therefore be mediated by diversity in co-factors, organizations of binding sites and conformational changes in the transcription factor itself. For example, human GCM1 gains a novel recognition sequence when paired with the ETS family factor ELK1 (*1*); auxin-responsive transcription factors regulate expression differentially depending on the arrangement of their binding sites (*2*); and the heat shock factor Hsf1 senses increased temperature by changing its conformation, which allows it to bind a unique recognition sequence (*3*).

We sought to investigate the features of a single transcription factor that uses distinct co-factors, binding sites and environmental inputs to mediate a cellular decision. The Ste12 protein of the yeast *Saccharomyces cerevisiae* governs the choice of mating or invasion. Each of these traits contributes to cellular fitness: mating of haploid yeast cells is required for meiotic recombination, and invasion allows a cell to forage for nutrients or to penetrate tissues, a characteristic of pathogenic fungi. Mating is initiated by binding of the appropriate pheromone, which activates an evolutionarily conserved G-protein-coupled MAPK pathway (*4*, *5*) (Fig. 1A). By contrast, invasion is initiated in response to increased temperature and limited nutrient availability (*6*–*8*). The shared MAPKKK Ste11, a client of the chaperone Hsp90 (*9*), likely contributes to the environmental sensitivity of both traits.

**Figure 1.**
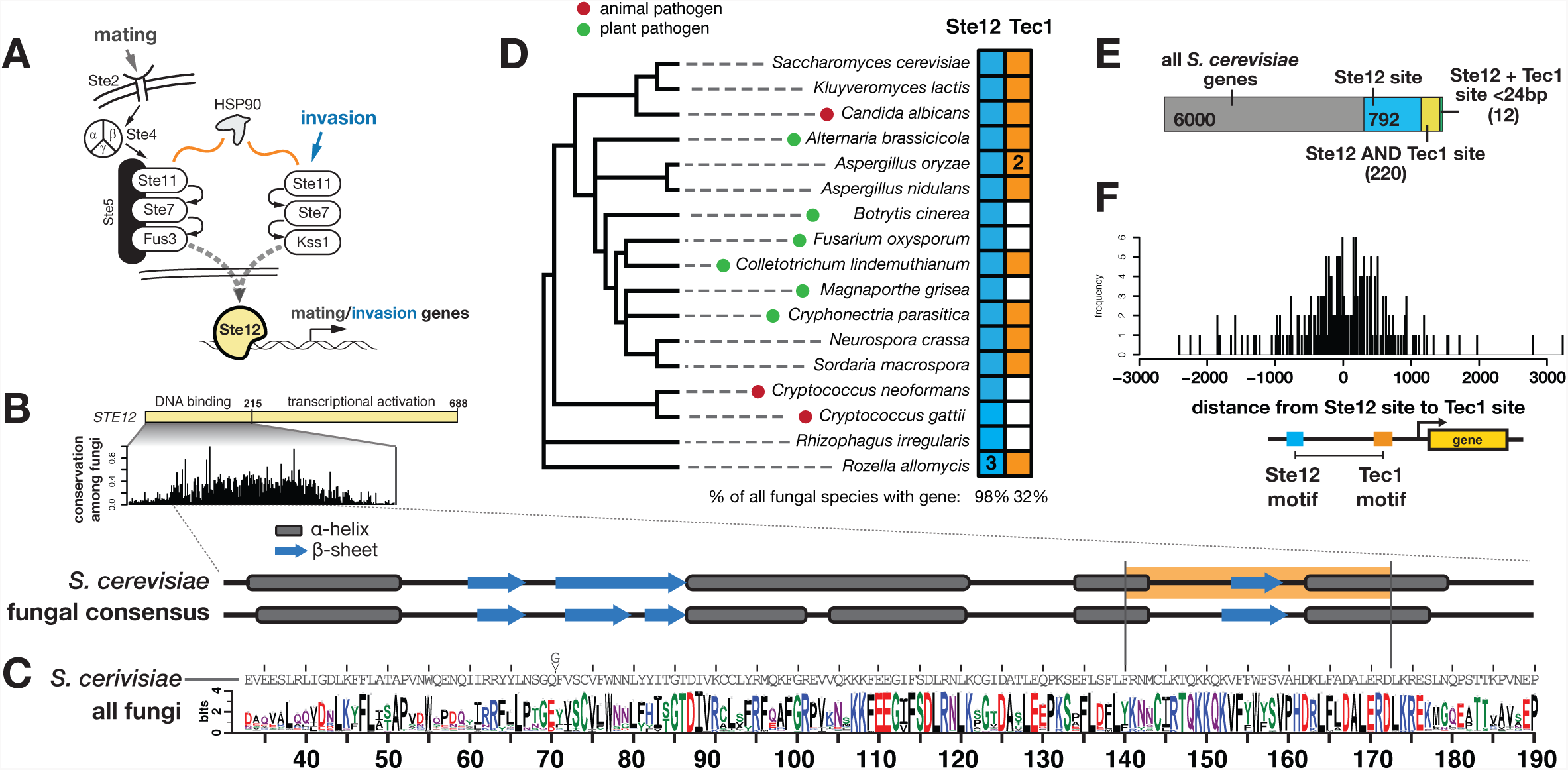
Conservation of Ste12 and Tec1 and their DNA-binding sites. (A) The yeast mating and invasion pathways contain shared signaling components, and both depend on Ste12 for activation of distinct regulatory programs. (B) *S. cerevisiae* Ste12 protein has a non-conserved transcriptional activation domain, as well as a highly conserved DNA-binding domain. The secondary structure of this domain is predicted to contain a pattern of alpha-helices (grey boxes) interspersed with β-sheets (blue arrows) that are conserved among all fungal species. The region of the DNA-binding domain chosen for mutagenesis is shaded in orange. (C) A logo plot of the conserved portion of Ste12’s DNA-binding domain generated from 1229 fungal species. (D) A phylogeny of fungal species selected for those with functionally tested *STE12* genes (except for the basal fungi *R. irregularis* and *R. allomycis* shown as outgroups). Pathogenic species are indicated with circles, and colored according to plant (green) or animal (red) hosts. All fungal species were queried for presence of a *STE12* (blue) or *TEC1* (orange) gene, and filled squares indicate presence of either gene, numbers inside boxes indicate species with multiple gene copies. Results for all species are shown below each gene’s column. (E) Proportional bar chart showing the number of *S. cerevisiae* genes that contain Ste12 sites (blue), Tec1 and Ste12 sites (yellow), and adjacent sites less than 24 bases from each other (green). (F) Histogram showing frequencies of spacing between all Ste12 and Tec1 binding sites in the *S. cerevisiae* genome. Negative values indicate the Tec1 site is 5’ of the Ste12 site, and positive values the converse.

Specificity between mating and invasion pathways is generated at multiple levels. For example, a mating-specific scaffold protein, Ste5, guides kinase signal transduction to activate mating (*10*). Two MAP kinases, Fus3 and Kss1, have overlapping functions in mating but opposing functions in invasion (*5*). The two pathways converge on Ste12, which interacts differentially with cofactors to activate either mating or invasion. For mating, Ste12 can bind at pheromone-responsive genes as a homodimer or with the cofactors Mcm1 or Mata1 (*11*–*13*). The consensus DNA-binding site of Ste12 is TGAAAC, known as the pheromone response element (PRE) (*11*). For invasion, Ste12 and its co-factor Tec1 are both required to activate genes that mediate filamentation (*8*, *14*, *15*). Some of these genes contain a Ste12 binding site near a Tec1 consensus sequence (TCS) of GAATGT, an organization for heterodimeric binding known as a filamentation response element (FRE). An alternative model, however, posits that expression of invasion genes is driven by a complex of Ste12 and Tec1, acting solely through Tec1 binding sites (*15*). An environmental component of trait specificity has been shown in fungi that are animal pathogens; upon recognition of host body temperature (37°C) (*6*), *Cryptococcus neoformans* has decreased mating efficiency, but increased ability to invade (*16*). Although trait preference in *S. cerevisiae* depends on Ste12, its role in regulating mating and invasion in response to increased temperature is unknown.

Because the highly conserved Ste12 DNA-binding domain ultimately enacts the choice between mating and invasion, here we sought to precisely define its DNA-binding specificity and examine the consequences of mutations and increased temperature on the balance of these traits. We reveal distinct organizational preferences for both homodimeric binding sites and cooperative heterodimeric binding sites with Tec1. Further, single amino acid changes suffice to shift the preference of yeast cells toward one or the other trait; they also suffice to confer dependence on the chaperone Hsp90 and responsiveness to higher temperature. Some of the hyperinvasive separation-of-function mutations are independent of the co-factor Tec1, thought to be essential for invasion. We show that these Ste12 variants bind to homodimeric Ste12 binding sites in which one site is highly degenerate. Such a binding preference provides a plausible mechanism to activate the expression of invasion genes in a Tec1-independent manner, and provides a model for the regulation of these genes in the many fungal species that have no copy of *TEC1*.

## Results

### *STE12* is present in nearly all fungal genomes, but most lack a *TEC1* orthologue

As the target of two different MAPK signaling cascades, Ste12 acts to define pathway specificity at the level of transcription (Fig. 1A). This role in both mating and invasion appears to be conserved in other fungal species, and presumably derives from properties of its DNA-binding domain, the most conserved segment of the protein (Fig. 1B) (*17*). Although the Ste12 DNA-binding domain has no match outside of the fungal kingdom, several residues as well as the overall predicted secondary structure are conserved in fungi (Fig. 1B, 1C). We addressed the cooccurrence of Ste12 and Tec1 by examining 1229 fungal species for the presence or absence of these two genes. Nearly every fungal species contains a copy of *STE12* (97.7% of species), while fewer than one third (31.7%) appear to have a copy of *TEC1* (Fig 1 - Suppl. Fig. 1). Furthermore, even among fungal pathogens with a characterized role for Ste12 in invasion, we found several examples in which a *TEC1* gene is not present (Fig. 1D). Therefore, fungal species must have evolved Tec1-independent strategies to regulate mating and invasion.

We tested the extent to which Ste12 and Tec1 binding at adjacent sites within filamentation response elements could explain invasion-specific activation by Ste12 DNA in *S. cerevisiae*. Cooperative binding at adjacent (within 24 bp) Ste12 and Tec1 binding sites (FREs) has been demonstrated *in vivo* and *in vitro* (*18*), although most invasion genes do not contain FREs (*15*). For all promoter sequences in *S. cerevisiae*, we assessed the frequency and spacing of sites, and called adjacent sites (<24 bp). We found 792 unique promoters with matches (*p* < 1e^−4^) to a Ste12 binding site, with 220 (27.8%) of these also having a Tec1 binding site (Fig. 1E). The median distance between Ste12 and Tec1 sites is 360 base pairs, and only twelve pairs of sites (5%) are within 24 base pairs of each other (Fig. 1F). Among these twelve, four have overlapping motifs consisting of a tail-to-tail arrangement, with the Tec1 motif overlapping the 3′ end of the Ste12 motif (Fig. 1F); this organization is more likely than non-overlapping sites to occur by chance and may not be functional, though overlapping sites have been observed in cooperative binding of mammalian transcription factors (*1*). The small number of sites organized for cooperative binding with Tec1 contrasts with the hundreds of genes upregulated under invasion conditions (*19*, *20*) as well as with the dozens of genes bound by Ste12 under invasion conditions (57 genes unique to invasion, 100 overall) (*14*). Thus, the model of Ste12 and Tec1 cooperatively binding to FREs within invasion genes cannot alone account for the broad transcriptional response during invasion observed in *S. cerevisiae*. Specificity may derive from instances of long-range looping interactions, or binding of Ste12 to DNA indirectly through its interaction with Tec1 bound at Tec1 consensus sequences (*15*). However, given the absence of Tec1 in many species, this model is also unlikely to be applicable in fungi more broadly.

### Ste12 and Tec1 preferentially bind DNA in defined spacings and orientations

Because the yeast genome contains few filamentation response elements, we sought to determine at high-resolution the *in vitro* DNA-binding preferences of Ste12 and Tec1. We used high-throughput systematic evolution of ligands by exponential enrichment (HT-SELEX) (*1*, *21*, *22*) with a random sequence of 36 base pairs, which allowed us to capture instances of Ste12 and Tec1 binding sites in all orientations and at every possible spacing of 24 base pairs or fewer. In the Ste12 sample, we captured monomeric or homodimeric instances of the expected Ste12 binding site of TGAAAC, with dimeric sites found 20-fold more frequently than monomeric sites (Fig. 2 - Suppl. Fig. 1C). Many transcription factors exhibit strong preferences for spacing and orientation of sites, and this preference is conserved within transcription factor families (*1*, *22*). Ste12 showed a strong preference (33% of 2,229,168 bound output sequences) for tail-to-tail binding sites with a three base pair spacer (Fig. 2A, top). While the most enriched sequences contained two perfect TGAAAC sites with this spacing (Fig. 2B, top), most of these sequences contained one perfect TGAAAC site paired with a mismatched site. This strong spacing preference had not been identified in previous protein binding microarray studies, which used shorter target sequences (*23*), but is consistent with the known binding of Ste12 as a homodimer (*24*). We found two instances of homodimeric sites with a perfect TGAAAC pair in the *S. cerevisiae* genome: in the promoters of *GPA1*, encoding the α-subunit of the G protein involved in pheromone response, and of *STE12* itself (Fig. 2C). Gpa1 and Ste12 are a component of the initial signaling point after pheromone sensing and the ultimate transcription target of the MAPK signaling cascade, respectively, and both are among the most highly induced genes in response to pheromone (*25*). Elsewhere in the genome, three base pair-spaced sites in which one site is perfect and one is mismatched are found within several known pheromone-regulated genes. This configuration occurs in 29 genes in the *S. cerevisiae* genome, and 12 of the 50 most pheromone induced genes contain these sites (examples in Fig. 2C). To analyze the ability of these sites to drive Ste12-dependent expression, we used a reporter assay in yeast. The *in vitro* preferences for the dimeric sites, as well as for the base flanking the TGAAAC motifs, correlated with *in vivo* activation (Fig. 2 – Suppl. Fig. 1A, 1B).

**Figure 2.**
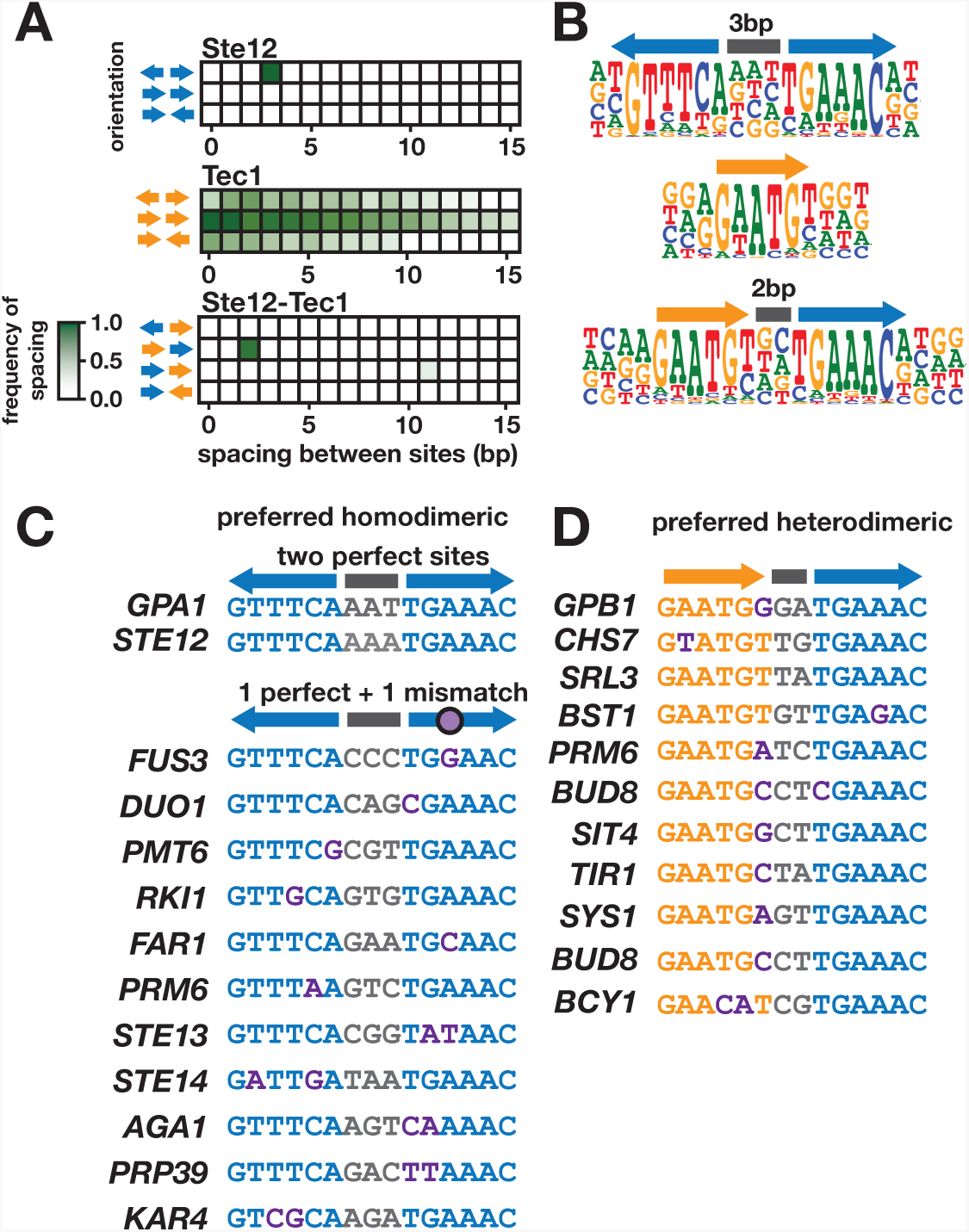
Identification of the DNA-binding preferences of Ste12 and Tec1 by HT-SELEX. (A) Heatmap showing relative frequencies of the possible orientations and spacings of the primary 6-mer selected in Ste12 (TGAAAC, upper) and Tec1 (GAATGT, middle) binding reactions. The bottom heatmap shows the frequency of each respective 6-mer in the co-binding sample containing both proteins. The single dark green box in the upper and bottom heatmaps indicates the most frequent orientation and spacing of sites; white boxes are at most 30% as frequent as the maximum. No frequent dimeric site organizations were observed for Tec1 (middle heat map). (B) Full motifs identified by Autoseed software (1). (C) Instances of the Ste12 homodimeric tail-to-tail, 3 base-pair spaced motif were used to query native yeast promoters. Two pheromone-induced genes, *GPA1* and *STE12*, contain two perfect sites, while most other genes contain a perfect site paired with a site containing one or two mismatches, with the 3 base-pair spacing intact. (D) A similar search of yeast promoters using the preferred Ste12 and Tec1 heterodimeric site identified a set of invasion-associated genes, as well as those not previously linked to invasion.

HT-SELEX with Tec1 protein recapitulated the known binding site of (A/G)GAATGT (Fig. 2B, middle panel) (*26*). We found that the first base of this motif showed a strong preference for a purine base, and that the final T showed less specificity than the rest of the motif. Unlike with Ste12, we did not observe any spacing and orientation preferences for two Tec1 sites, indicating that bound sequences with multiple Tec1 binding sites are likely the result of independent binding events (Fig. 2A, middle panel).

We next conducted HT-SELEX in the presence of both Ste12 and Tec1 to identify patterns of cooperative binding events between the two proteins. The DNA-binding domain of Ste12 (1-215) is sufficient for cooperative binding with Tec1 (1-280) *in vitro* (*26*, *27*). We detected bound sequences containing sites for both proteins, with the highest abundance preference being a tail-to-head orientation of Tec1 and Ste12 sites separated by two base pairs (Fig. 2A, 2B, lower panel; Fig. 2 - Suppl. Fig. 2.). Several known invasion genes in the *S. cerevisiae* genome contain this type of heterodimeric site, though some other genes with this binding site organization have not been previously associated with invasion (Fig. 2D). This arrangement is similar to homodimeric Ste12 sites, except that one Ste12 site is replaced by a Tec1 site, indicating a similar protein interaction surface may be used for this heterodimeric binding mode of Ste12 as is used for the homodimeric binding mode. However, even in this sample, the pool of bound sequences was dominated by two Ste12 sites with three base pair spacing, suggesting that Ste12 has higher affinity to two PRE sites than to an FRE requiring binding with Tec1.

### Mutations in the DNA-binding domain of Ste12 separate mating and invasion functions

The organization of DNA-binding sites selected by Ste12 alone or in combination with Tec1 suggested that Ste12 balances the expression of mating and invasion genes by its mode of binding to DNA. We sought to determine whether this balance could be shifted by mutations within the Ste12 DNA-binding domain. We conducted deep mutational scanning (*28*) of a segment of this domain by generating ~20,000 protein variants over 33 amino acids, including single, double, and higher order mutants, and subjected yeast cells carrying this variant library to selection for either mating or invasion (Fig. 3A; Fig. 3 – Suppl. Fig. 1; Fig. 3 – Suppl. Fig. 2A, 2B). As a control, we employed selection for a third trait, response to osmotic stress, which shares upstream pathway components but does not involve Ste12. Mating selections and osmotic stress selections were carried out in the BY4741 strain background. However, as this strain contains a *flo8* mutation that prevents invasion, we carried out invasion selections in a related strain, Sigma1278b (*29*), which has been used for large-scale invasion phenotyping (*30*). Although Sigma1278b and BY4741 have many genetic differences, Ste12 and Tec1 and their essential roles in balancing mating and invasion are conserved between the two strains.

**Figure 3.**
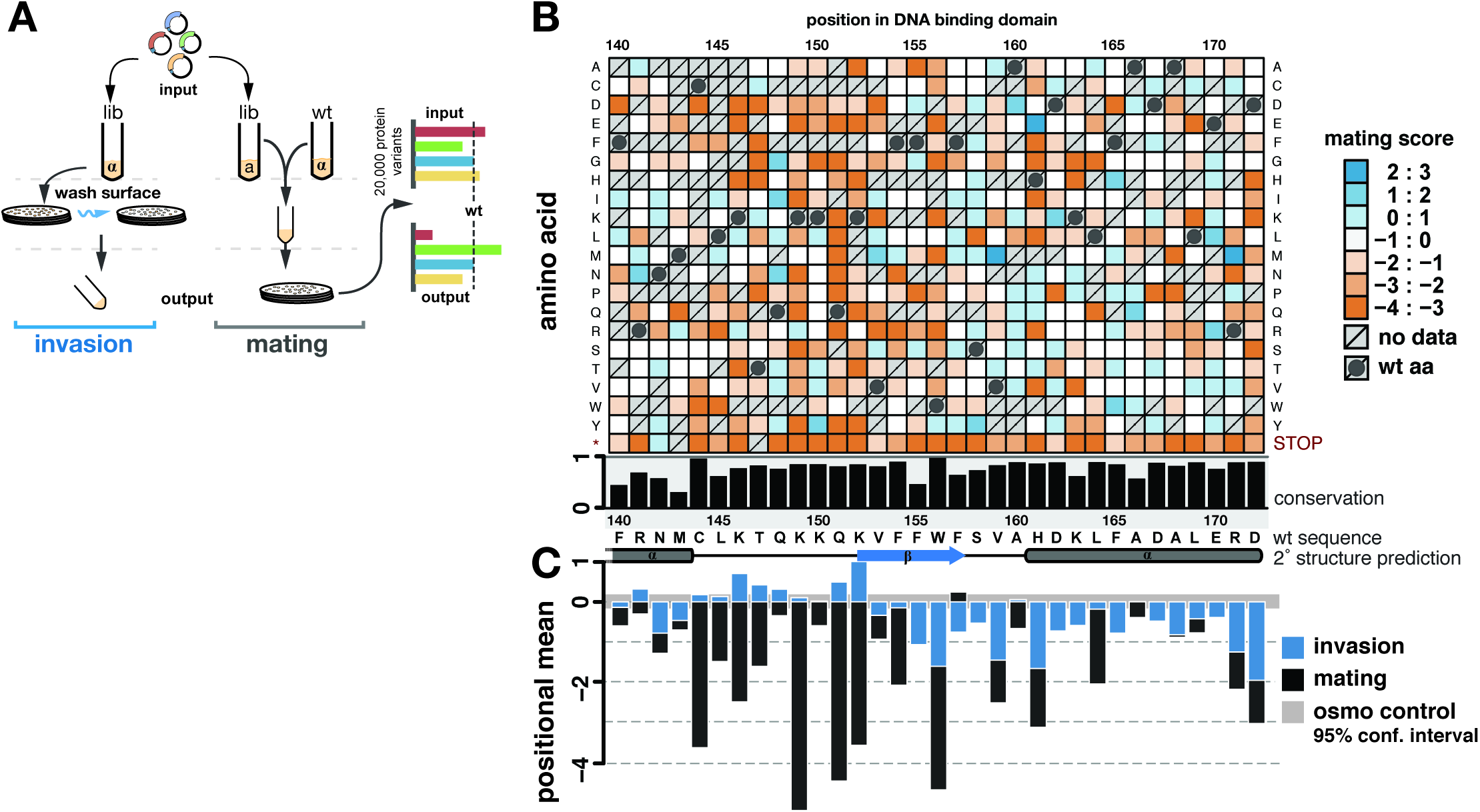
Deep mutational scanning identifies Ste12 DNA-binding domain mutants with altered mating and invasion function. (A) Two yeast populations were transformed with the same *STE12* variant library. For assaying invasion, SIGMA1278b *MATα* cells (n = 400,000 per replicate, 3 biological replicates) were plated on selective media, and grown. Selection was performed by washing colonies from plate surfaces and collecting cells embedded in the agar for sequencing. For assaying mating, BY4741 *MATα Ste12Δ* cells were mated to *MATα* cells, and diploids (n = 500,000 per replicate, 3 biological replicates) were selected, scraped from the agar, and sequenced. In both cases, input variant frequencies were defined prior to selection. (B) The effects of single amino acid substitutions within Ste12’s DNA-binding domain on mating ability are shown. On the x-axis, the wild-type Ste12 sequence is shown, along with its predicted secondary structure (helices shown as tubes) and conservation. Conservation was determined as fraction of identity among 1229 fungal species. On the y-axis, amino acid substitutions are shown. Variants increasing mating efficiency are in shades of blue, and variants decreasing mating efficiency are in shades of orange based on log2 enrichment scores relative to wild-type. Dark grey circles indicate the wild-type Ste12 residue, and crosses indicate missing data. Ste12 variants showed comparable expression levels (Fig. 3 - Suppl. 3). (C) Positional mean scores for single amino acid substitutions are shown for mating (black) and invasion (blue), excluding stop codons. Grey horizontal bar indicates confidence interval for experimental noise determined from selection for Ste12-independent high osmolarity growth.

We expected that most *STE12* mutations would affect mating and invasion similarly, because of the conservation of the Ste12 DNA-binding domain and its requirement for both traits. Indeed, we found positions in which almost any amino acid substitution was highly deleterious to both mating and invasion (Fig. 3B, 3C; Fig. 3 - Suppl. Fig. 2). We identified sites that were more sensitive or less sensitive, on average, to mutation, by calculating a positional mean score from all mutations tested at that site. Each mutation was tested in triplicate, and the experimental error was calculated for each mutant individually such that standard error of the positional mean represents the variability among different amino acid substitutions, rather than experimental noise (Fig. 3C, Fig. 3 – Suppl. Fig. 1E). Conservation only partially explained these results, as some of the most deleterious positions are invariant among fungi (W156, C144), whereas others are not (K149, Q151, K152). Differing expression levels among Ste12 variants were not predictive for either mating or invasion phenotypes (Fig. 3 – Suppl. Fig. 3).

The mutational analysis revealed “separation-of-function” mutations, which primarily reduced either mating or invasion. Substitutions with mostly deleterious effects on mating clustered in residues N-terminal to the conserved W156 (Fisher’s exact test p-value = 0.002, designated region I), while substitutions with increased effects on invasion appeared more frequently in this region (p-value = 0.0003). Substitutions that increased mating were rare. By contrast, substitutions at some positions, almost all within region I, both increased invasion and decreased mating. We verified separation-of-function mutants, and further explored the apparent structure of mutational effects with a much larger pool of 20,000 double mutants. Double mutants involving two positions in region I were mostly defective for mating, while those involving two positions in region II were mostly defective for invasion, confirming the bipartite arrangement of the mutagenized segment, split by the central W156 (p-value = 1.3e-5, Fig. 4A). Double mutants between regions I and II showed variability in mutational effects, with some pairwise combinations increasing mating and others decreasing mating; the effects on invasion, however, for these same combinations did not follow the same pattern, suggesting epistatic interactions (examples highlighted in Fig. 4A, Fig. 4 – Suppl. Fig. 1). Thus, mutations in the Ste12 DNA-binding domain can impose preference for mating or invasion rather than similarly affecting both traits, suggesting that the DNA-binding domain itself contributes to trait specificity.

**Figure 4.**
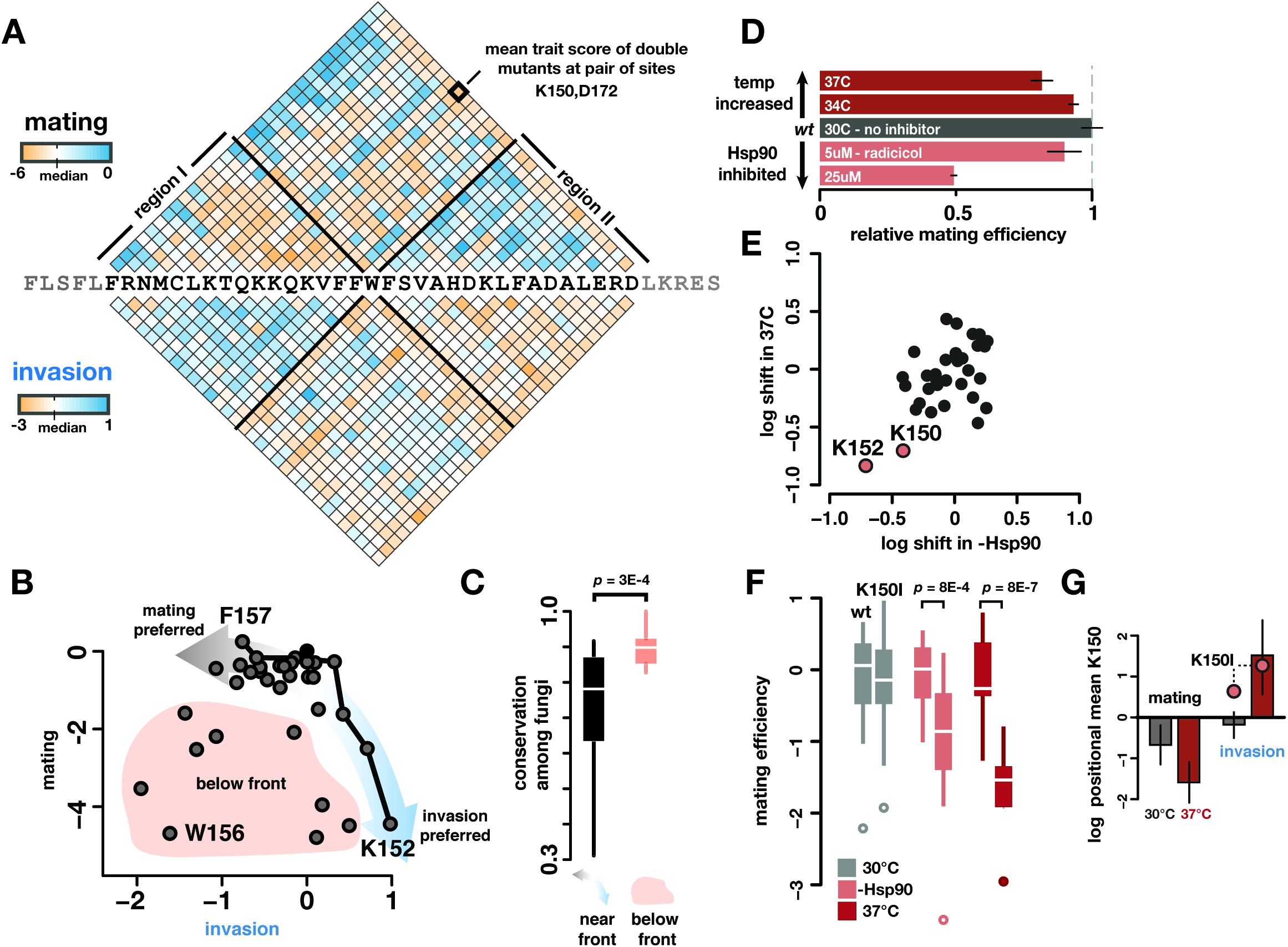
Mutations residing in a region of the Ste12 DNA-binding domain increase invasion at the cost of mating, with exceptional mutations doing so in a temperature-and Hsp90-dependent manner. (A) Mean effect of double mutations between all combinations of positions for mating (above wild-type sequence) and invasion (below wild-type sequence). A specific pair of positions is shown in bold. Effects are color-coded as in Figure 3, functional variants in shades blue, nonfunctional variants are shown in shades of orange. Note difference in scale ranges between mating and invasion. Black lines emanating from W156 represent boundaries of region I, in which mutations primarily reduced mating, and region II, in which mutations primarily reduced invasion. (B) Scatterplot of positional mean scores for both mating and invasion showed inverse relationship, indicating a tradeoff between both traits. The tradeoff is visualized as a Pareto front (black line), which was determined empirically; positions near the front maintain high values for one trait and minimize costs to the other. Arrows indicate preference for either mating (grey) or invasion (blue). Positions near the front were distinguished from those below the front (shaded in red) by calculating Euclidian distances. (C) Boxplots show conservation for positions near and below the front; positions below the front are significantly more conserved among fungi than those near the front (two-sided t-test). (D) Mating efficiency of yeast cells with wild type Ste12 at increased temperature (dark red) or in the presence of Hsp90 inhibitor radicicol (pale red) is reduced relative to an untreated control (black), error bars represent standard error of the mean (s.e.m). (E) The mating efficiency of Ste12 variants at high temperature or with Hsp90 inhibition is shown as the shift in mean effect at each mutated position. Two positions, K150 and K152 (pale red), showed greatest sensitivity to both treatments. (F) K150I was tested in a quantitative assay to validate its temperature-sensitive and Hsp90-dependent mating activity, shown relative to wild type (left boxplot in each pair) in each treatment (n = 20 for each sample). (G) Ste12 variants at K150 (mean effect) decreased mating (left panel) and invasion (right panel) at standard temperature (grey bars). At high temperature (red bars), mating further decreased; however, invasion increased. Error bars represent s.e.m. K150I variant is individually highlighted for comparison.

To explore the apparent tradeoff between mating and invasion in more detail, we used the Pareto front concept (*31*). This concept, rooted in engineering and economics, defines all feasible solutions for optimizing performance in two essential tasks. Here, we consider all feasible genotypes that affect mating or invasion. We plotted the single mutation mean positional scores for both traits, identifying positions close to the Pareto front that show increased invasion and deleterious effects on mating. (Fig. 4B, Fig. 4 – Suppl. Fig. 2). We find this effect is recapitulated in the effects of individual mutants tested in both trait selections at extreme positions on the front (Fig. 4 – Suppl. Fig. 3). Positions well below the Pareto front, including most prominently W156, had deleterious effects on both traits. These positions are significantly more conserved among fungi, consistent with variation at these sites being disfavored given the constraint on Ste12 to maintain both mating and invasion function in the fungal lineage (Fig. 4C). That opposing preference for each trait is better predicted by positional means than every individual mutation suggests that trait preference is encoded in particular functional regions of the Ste12 DNA-binding domain rather than through overall features like protein stability.

Having established that trait preference can be modulated by mutations in Ste12, we asked whether temperature affected specificity toward mating and invasion, and if that specificity could be altered by mutation. Furthermore, since chaperones maintain protein function at increased temperatures and Hsp90 modulates pheromone signaling in *S. cerevisiae* (*9*), we also asked whether mating or invasion changed in the presence of radicicol, a pharmacological inhibitor of Hsp90. We confirmed that mating is modulated by temperature and by Hsp90 function by mating cells expressing wild-type Ste12 at increased temperature or in the presence of radicicol (Fig. 4D). We then subjected the Ste12 variant library to mating selection at 37°C or in the presence of radicicol. Most Ste12 variants responded to increased temperature or Hsp90 inhibition as did wild-type Ste12 (Fig. 4 – Suppl. Fig. 5). However, there were two positions, K150 and K152, in which mutations resulted in highly temperature-responsive and Hsp90-dependent mating (Fig. 4E). This pair of lysines resides within the mutagenized segment that modulates mating and invasion specificity. We validated the temperature and Hsp90 effect on a variant (K150I) that conferred mating at near wild-type levels in the absence of heat or radicicol treatment (Fig. 4F). However, in the presence of heat or radicicol treatment, mating of cells with the K150I variant was severely decreased, indicating that this mutation led to buffering by Hsp90. Thus, Hsp90 could facilitate a mutational path toward the pathogenic lifestyle by minimizing mating costs at 30°C and enhancing invasion at 37°C. To test this idea, we conducted selection for invasion at 37°C, which yielded results comparable to Hsp90 inhibition for mating. Indeed, the mean effect of all variants at K150 was to increase invasion at high temperature with a concomitant decrease in mating, and this was also true of the individually validated variant K150I (Fig. 4G), demonstrating that K150 variants were not simply unstable at high temperature but gained a novel function. Thus, mutations in Ste12 can interact with an environmental factor to further bias cellular decision-making toward invasion over mating. Mutation to Ste12 thereby allows *S. cerevisiae* to mimic the behavior of fungal pathogens like *Cryptococcus neoformans*, for which the sensing of the increased body temperature of their animal hosts facilitates the transition toward an invasive lifestyle (*7*).

**Figure 5.**
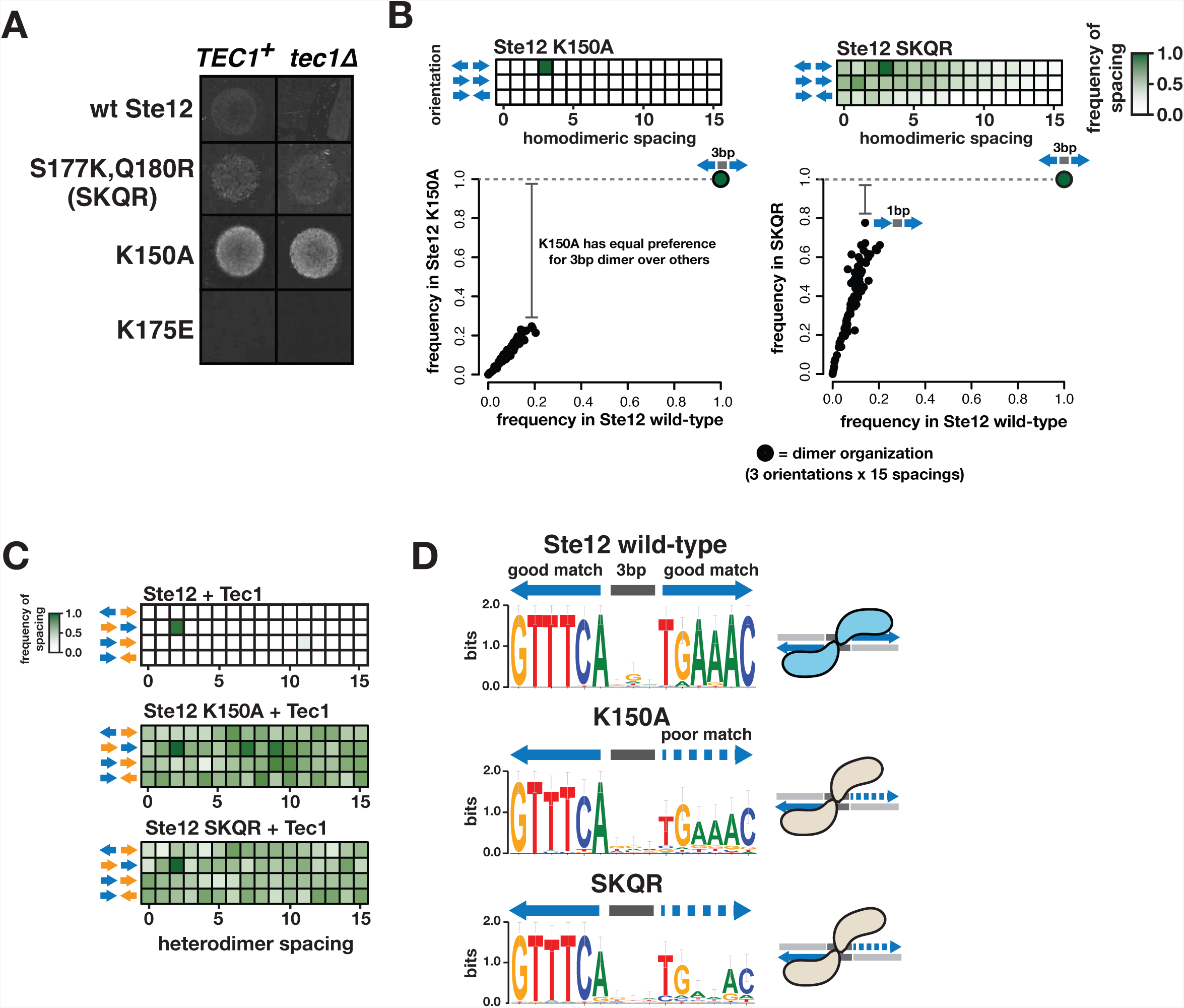
Ste12 DNA-binding domain mutants that confer invasion independent of the co-factor Tec1 have altered binding specificity. (A) Wild-type Ste12 requires the co-factor Tec1 for invasion; a *tec1Δ* strain fails to invade. The region I mutation K150A showed a hyperinvasive phenotype with or without Tec1. Introduction of the two positively charged residues found in *C. gattii* (SKQR) also led to Tec1-independent invasion, whereas introduction of a negative charge (K174E) eliminated invasion. The hyperinvasive Ste12 mutants showed a dominant invasion phenotype, as they were tested in the presence of wild-type Ste12, in the SIGMA1278b background. (B) Spacing heatmaps for Ste12’s binding site are shown as in Fig. 2A. The K150A retains the same preference for tail-to-tail 3 base pair-spaced sites as the wild-type Ste12, but SKQR’s preference is reduced. To visualize this difference, the scatterplots below show relative frequencies of site organizations for each variant (y-axis) compared to wild-type Ste12 (x-axis), where each point represents one orientation and spacing combination (example of SKQR’s reduced preference is shown: its preference for head-to-tail, 1bp is ~78% that of the tail-to-tail 3bp site). (C) Spacing heatmaps are shown for the K150A and SKQR variants in co-binding reactions with Tec1; unlike wild-type Ste12 (repeated from Fig. 2A lower panel for clarity), neither variant shows strong preference for any heterodimeric site organization. (D) Logo plots generated from the 50 most highly enriched sequences in HT-SELEX represent ideal sites for each variant alongside wild-type Ste12.

We examined whether natural variation in fungal Ste12 DNA-binding domains includes amino acid residues found to affect *S. cerevisiae* trait preference, as fungal species have differential capacities for mating and invasion (*17*, *32*). We examined variation in the DNA-binding domain present in species with and without a *TEC1* gene (Fig. 5 – Suppl. Fig. 1) and detected variation outside of the mutagenized region that might be expected to benefit invasion. For example, the Ste12 DNA-binding domain of the invasive pathogen *Cryptococcus gattii* (*33*) contains two additional positive residues immediately C-terminal to region II; such changes are found in many other species that lack Tec1 (Fig. 5 – Suppl. Fig. 1). Introducing these *C. gattii*-specific residues (S177K, Q180R; denoted SKQR) into the *S. cerevisiae* Ste12 DNA-binding domain yielded a dominant invasion phenotype (Fig. 5A) and decreased mating (Fig. 4 – Suppl. Fig. 4) in *S. cerevisiae*. We introduced a negatively charged residue (K175E) into this same region and abolished invasion entirely (Fig. 5A). Similarly to SKQR, the region I mutation K150A conferred decreased mating and an increase in invasion. The mating defect and dominant invasion phenotype for each variant suggest that these Ste12 variants engage in altered binding to DNA, altered homodimerization, or altered heterodimerization with Tec1.

### Mutations in Ste12 can promote Tec1-independent invasion and alter DNA-binding specificity

Tec1 activates invasion genes, with Ste12 either binding directly to DNA cooperatively with Tec1 or indirectly as part of a complex with Tec1 (*14*, *15*). However, in contrast to both established Ste12 binding modes, the invasion phenotype due to the SKQR variant was independent of Tec1, as was the even stronger invasion phenotype due to the K150A variant (Fig. 5A). Thus, a single mutation in the Ste12 DNA-binding domain might change the binding preference of this domain such that it can be recruited in the absence of Tec1 to binding sites sufficient for invasion.

To understand the relationship between the phenotypes conferred by Ste12 variants and their DNA-binding specificity, we conducted HT-SELEX experiments. We chose the SKQR and K150A variants, each of which conferred a mating defect, and asked whether they could recognize the homodimeric Ste12 sites favored by the wild-type protein. SKQR showed a greatly reduced (by 45%) preference for two sites separated by three base pairs (Fig. 5B). Consistent with this reduced preference, SKQR conferred reduced activation *in vivo* from a minimal promoter containing such homodimeric sites (Fig. 5 - Suppl. Fig. 2A). Furthermore, an examination of the most enriched sequences bound *in vitro* by the SKQR variant showed that it lost specificity for the sequence of the second site; this was also reflected in its *in vivo* sequence preference (Fig. 5D, Fig 5 – Suppl. Fig. 5B). In contrast, K150A retained a strong preference for the three base-pair spaced sites, nearly identical to wild-type Ste12 (Fig. 5B). However, analysis of its most enriched sequences revealed that, like SKQR, the K150A variant showed reduced specificity for one site of the homodimeric pair (Fig. 5D). The reduced specificity in these variants should expand the number of their recognition sites. Because both of these variants recognized at least one perfect, or near perfect, Ste12 site, we conclude that the primary defect in these variants derives not from loss of specific DNA contacts, but rather from their diminished capacity for symmetric dimeric binding.

We also carried out co-binding SELEX experiments with Tec1, as previously conducted with wild-type Ste12 (reproduced in Figure 5C, top). Consistent with their ability to promote Tec1-independent invasion, both SKQR and K150A showed altered patterns of cooperative binding with Tec1. K150A showed no preference for the two base pair-spaced heterodimeric sites favored by the wild-type Ste12 in co-binding experiments with Tec1, and SKQR had a greatly reduced preference for these motifs (Figure 5C).

## Discussion

Fungal species make the decision to mate or invade based on environmental cues such as temperature, with the choice enacted through trait-specific transcription factors (*8*). In *S. cerevisiae*, the highly conserved Ste12 protein drives expression of mating genes as a homodimer and of invasion genes as a heterodimer with the invasion co-factor Tec1; the two proteins bind cooperatively to DNA (*18*). However, few invasion genes contain Ste12 and Tec1 binding sites spaced closely enough to plausibly allow efficient heterodimeric binding as the basis of trait specificity. One model that explains the scarcity of cooperative binding sites is based on the finding that Tec1 alone can bind to invasion genes, which then recruits Ste12 to provide activation activity without Ste12 itself binding DNA (*15*). However, only about a third of 1229 sequenced fungi contain a *TEC1* orthologue; moreover, the vast majority of these fungi contain only a single *STE12* gene, excluding *STE12* gene duplication and subsequent sub-functionalization as a prominent mechanism of achieving trait specificity.

Here, we show that single mutations in Ste12 are sufficient to shift yeast trait preference toward invasion. One possible explanation for how such mutations separately affect the two traits is that they alter the interaction of Ste12 with Tec1. However, this explanation cannot be valid, as the hyperinvasive Ste12 variants were independent of Tec1 presence. This Tec1-independent phenotype provides a plausible mechanism for how fungal species without Tec1 or similar co-factors accomplish activation of invasion genes. Indeed, the S177K change in the Ste12 of *C. gatti* causes a hyperinvasive phenotype when introduced into the Ste12 of *S. cerevisiae* and is more frequently found in fungal species without Tec1 (Fig. 5 - Suppl. Fig. 1). In contrast, there is little natural variation in the positions of Ste12 in which mutations shift trait preference towards invasion and change the response to temperature and Hsp90 inhibition. Such conservation suggests that most fungal species have retained the capacity for both invasion and mating, even though many have not yet been found as diploids. Further, our finding of dominant hyperinvasive Ste12 variants points to an explanation for the near universal absence of Ste12 duplication and sub-functionalization: Ste12 paralogs optimized for invasion in this manner would likely suppress mating.

Just as single mutations could eliminate Ste12 dependence on Tec1 for driving gene expression, they could also change the response of Ste12 to temperature. Increased temperature, reflecting the body temperature of animal hosts, can promote invasion of fungal pathogens (*7*). Even in the non-pathogenic *S. cerevisiae*, high temperature decreases mating and promotes invasion. Mutations at the same positions in Ste12 implicated in shifting trait preference could also alter dependence on temperature and the chaperone Hsp90. The temperature-and Hsp90-dependent variants did not resemble typical temperature-sensitive mutants in simply losing function under non-permissive conditions; in response to high temperature or Hsp90 inhibition, these variants failed to promote mating yet conferred an increased ability to invade. These variants thus represent the unusual phenomenon whereby perturbation of Hsp90 or equivalent environmental stress results in a protein gaining a novel function at the cost of another, whereas more typically, mutated proteins (such as many oncogenic kinases) are enabled to function by Hsp90 and fail to do so in response to high temperature or Hsp90 inhibition (*34*, *35*).

Although Hsp90 broadly buffers the phenotypic consequences of genetic variation, only a handful of Hsp90-dependent variants have been mapped to date, limiting our understanding of their prevalence and biological significance (*36*, *37*). As the consequences of human disease mutations increasingly are found to be dependent on Hsp90, deep mutational scanning provides a possible experimental avenue to systematically identify features of Hsp90-dependent variation (*38*, *39*). In Ste12, such mutations are rare and position-dependent, consistent with their effects on protein folding. Yet the Hsp90-dependent Ste12 variants are accessible via a single mutation, suggesting that the chaperone could facilitate a mutational path toward a pathogenic fungal lifestyle by minimizing mating costs at normal temperature and enhancing invasion at higher host temperature.

The ease with which Ste12 can shift trait preference towards invasion – independent of Tec1 and modulated by Hsp90 and temperature – poses the question of how this shift is achieved at the level of DNA binding. We show that wild-type Ste12 strongly preferred two tail-to-tail binding sites separated by three base pairs, consistent with its binding as a homodimer. The majority of homodimeric Ste12 sites in the *S. cerevisiae* genome are organized in this way, containing a perfect site paired with a site carrying one or more mismatches.

Hyperinvasive Ste12 variants also bound to dimeric sequences, but while one was a perfect or near-perfect Ste12 site, the other was highly degenerate. Addition of Tec1 did not yield cooperative binding sites, consistent with the Tec1-independence of these variant Ste12 proteins *in vivo*. Since many invasion genes contain perfect or near-perfect Ste12 binding sites adjacent to highly degenerate ones, the binding preference of the hyperinvasive variants provides an explanation for how they can enact the invasion program in the absence of Tec1. However, the same variants failed to activate expression of mating genes, which tend to contain two nearly perfect sites, as well as the wild-type Ste12 protein did.

We propose the following model to explain these data (Figure 6). In mating genes, two properly oriented binding sites facilitate the interaction of two Ste12 proteins. Under invasion conditions, the Ste12 dimer interface instead allows cooperative interaction between Ste12 and Tec1. We posit that mutations in Ste12 that lead to hyperinvasiveness change this interface such that the spatial conformation of the homodimer is altered and interaction with Tec1 is not possible. In this altered conformation, the Ste12 proteins, unable to make a proper symmetric contact, lose their preference for the perfect/near-perfect homodimeric binding sites. However, the altered conformation confers the ability for Ste12 to recognize a much more degenerate version of the binding site adjacent to a near-perfect site; likely neither site is bound as tightly as the wild-type Ste12 protein binds to canonical sites. The proposal of an altered Ste12 protein conformation is supported by the finding of temperature-and Hsp90-dependent variants in the Ste12 region that mediates the shift towards invasion.

**Figure 6.**
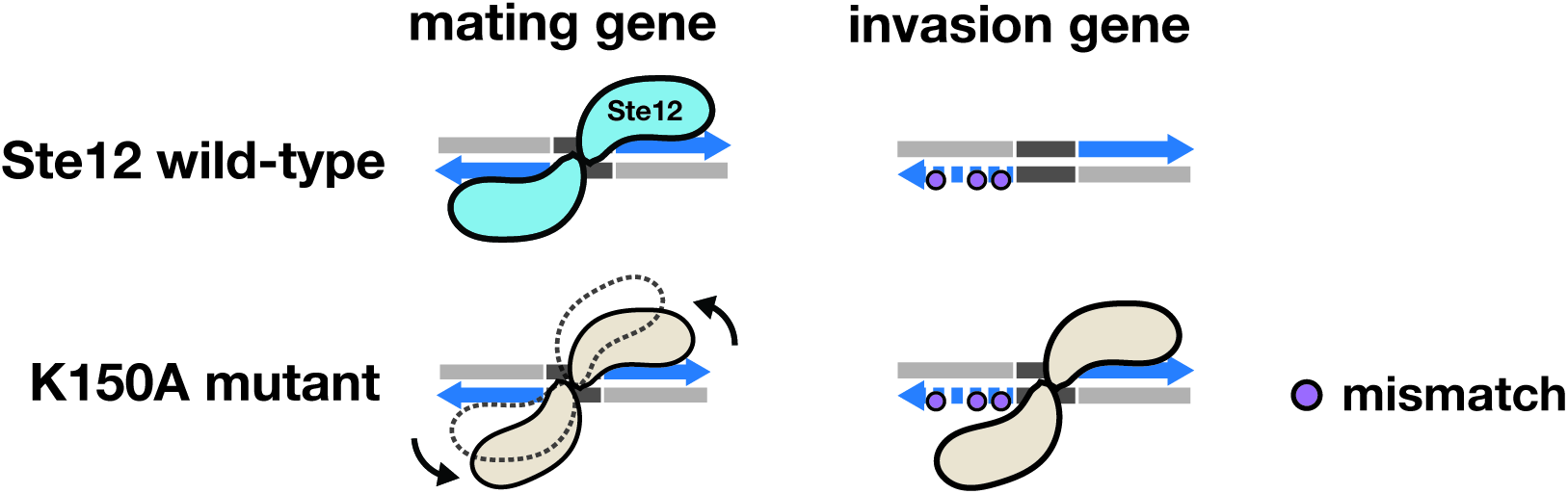
A model for novel site recognition by Tec1-independent Ste12 variants. Wild-type Ste12 protein (blue) is compared to the K150A mutant (tan) for binding at a mating gene with two perfect recognition sites (tail-to-tail with a 3 base pair spacing, shown in blue) and an invasion gene with a single perfect site along with a degenerate site. Due to its altered dimer interface, the “kinked” mutant K150A is less capable of binding symmetrically at two perfect sites, consistent with its phenotype of reduced mating efficiency. However, the conformation of K150A allows the variant protein to occupy novel sites within invasion genes that contain one perfect or nearly perfect site paired with a degenerate site; wild-type Ste12 is unable to occupy these sites.

Mutations in transcription factors that generate alternative DNA-binding modes have been previously identified. For example, rare coding variants in the homeodomain recognition helices of several transcription factors alter DNA-binding specificity (*40*), which likely contributes to associated diseases. As we found with the Ste12 variants, these homeodomain variants bind at sites not dramatically different from the wild-type sites, yet their promiscuity is associated with diseases. Expanding beyond these studies, we demonstrate here that Ste12 can explore alternative binding modes through both mutation and environmental stress, and thereby shift preference between two ordinarily mutually exclusive traits. This surprising malleability of transcription factor binding stands in stark contrast to the fact that sequences of transcription factors tend to be highly conserved. Indeed, throughout fungal evolution, the Ste12 DNA-binding domain remains largely invariant, with Ste12 taking advantage of both multiple interacting partners that have been gained and lost over time, as well as environmental conditions such as temperature, to influence decision-making. This study, together with those revealing only weak purifying selection in functional regulatory regions, poses the challenge of reconciling flexible transcription factor activity and often highly variable regulatory regions with the longstanding observation that gene expression patterns are robust to most perturbations and conserved throughout evolution.

## Materials and Methods

### HT-SELEX

We expressed and purified fragments of STE12 (1-215) and TEC(1 1-250) cloned into pGEX-4T-2 vectors. These protein fragments have been used previously and are sufficient for both individual cooperative binding *in vitro*. Fragments of each protein, as well as protein variants, were purified using a GST tag and subsequently used for HT-SELEX. SELEX reactions with homogenous and mixed protein populations were performed identically to previous work (*1*, *21*). Briefly, a 50uL reaction containing purified Ste12 and Tec1 (1:25 molar ratio with DNA), 200ng non-specific competitor double-stranded nucleic acid poly (dI/dC), 100ng selection ligand (36N) was incubated in binding buffer (140 mM KCl, 5 mM NaCl, 1 mM K_2_HPO_4_, 2 mM MgSO_4_, 20 mM HEPES [pH 7.05], 100 μM EGTA, 1 μM ZnSO_4_) for 2 hours. GST Sepharose (GE) beads were then added to each reaction, incubated for 30 minutes, and unbound ligand was removed using seven buffer washes. Output reactions were amplified by PCR after each round, and these products were subsequently used to prep high-throughput sequencing libraries.

### Generation of *STE12* mutant libraries

The *STE12* locus from *Saccharomyces cerevisiae* strain BY4741, including the intergenic regions, was introduced into the yeast vector pRS415 containing a *LEU2* marker (*41*). Degenerate DNA sequence encoding a 33 amino acid (99bp) segment of Ste12’s DNA binding domain was generated by 2.5% doped oligonucleotide synthesis (Trilink Biotechnologies, San Diego, CA). Invariant 30bp sequences were designed on either side of the mutagenized fragment; these flanking sequences contained NotI and ApaI cut sites found in the coding sequence of *STE12*, and were unique in the *STE12* plasmid construct. Both fragment and plasmid were double-digested with NotI and ApaI, and the plasmid library was assembled by standard ligation. *STE12* libraries were transformed into electrocompetent *E. coli* (ElectroMAX DH10B, Invitrogen), and amplified overnight in selective media. Efficiency of ligation was verified by Sanger sequencing across the mutagenized region of 96 transformants (*42*); no assembly errors were detected, and mutant proportions reflected those expected for doped oligo synthesis at 2.5%. Plasmid libraries were used to transform yeast (BY4741 MATa or Σ1278b-α) with a deleted endogenous copy of *STE12* by high-efficiency lithium acetate transformation (*43*). The same plasmid library was used to transform yeast (Σ1278b-α) for invasion selection. Individual point mutations were generated in wild-type *STE12* plasmids by site-directed mutagenesis (Q5, New England Biolabs).

### Large-scale trait selections

The BY4742 MATα strain was used as the mating partner for the library-transformed BY4741 MATa in all selections and mating assays. Transformed yeast cells were grown to late log-phase in a single 500mL culture, and cells were harvested to determine plasmid variant frequencies in the input population. The same culture was used to seed 36 independent mating selections for each treatment: one million MATa cells with *STE12* variants were mixed with 10-fold excess wild-type MATα cells and allowed 5 hours to mate(*44*). Depending on treatment type, cell mixtures were left at 30C with DMSO, 30°C with the Hsp90 inhibitor radicicol, or 37°C with DMSO. A 5uM concentration of the Hsp90 inhibitor radicicol (Sigma-Aldrich, R2146) was chosen due to its measureable effect on mating efficiency and lack of pleiotropic growth defects. Radicicol was chosen over the Hsp90 inhibitor geldanamycin (Sigma-Aldrich, G3381), because five-fold higher concentrations of geldanamycin were required to achieve the same phenotypic effect as with radicicol (data not shown). The temperature of 37°C was chosen for the similarity of effects on mating between temperature and radicicol treatments. After mating was completed, cell mixtures were plated using auxotrophic markers present only in mated diploids. Plasmids containing *STE12* variants were extracted from this output population, as well as from the pre-mating input, for subsequent deep sequencing. Mating selections were repeated in triplicate. For invasion, Σ1278b-α yeast cells transformed with the Ste12 plasmid library were grown to late log-phase in a single 500mL culture, diluted (10,000 cells per plate), and plated onto 40 plates of synthetic complete medium lacking leucine (2% agar). Plates were incubated at 30°C or 37°C for 72 hours to allow for sufficient invasion, as previously described (*30*). After three days, cells were washed from the plate surfaces, enriching for cells embedded in the agar. Agar pucks were removed from plates with a razor. Using the “salsa” blender setting (Hamilton Beach, Glen Allen, VA), a coarse slurry was generated and subsequently poured over a vacuum apparatus lined with cheesecloth. The resulting liquid cell suspension was spun down at 5000rpm to collect cells for subsequent deep sequencing. Individual invasion assays (as in Fig. 5A) were treated identically, but 10uL aliquots of OD-normalized cultures were plated to image colonies for each strain. For high osmolarity growth, selections were conducted using the BY4741 MATa library-transformed population grown overnight in media containing 1.5M Sorbitol, as described previously (*45*). Populations were sequenced before and after growth to determine enrichment scores that defined the 95% confidence interval used in Fig. 3C. For all trait selections, we chose sample sizes that were at least 10-fold higher than the variant library size, ensuring that each variant would be adequately sampled in each of the three biological replicate selections.

### Sequencing and determination of trait scores

Sequencing was completed on Illumina’s MiSeq or NextSeq platforms. Sequencing libraries were prepared by extracting plasmids from yeast populations (Yeast Plasmid Miniprep II, Zymo Research, Irvine, CA) before and after selection. These plasmids were used as template for PCR amplification that added adaptor sequences and 8bp sample indexes to the 99bp mutagenized region for sequencing (all libraries amplified < 15 cycles). Paired-end reads spanning the mutagenized region were filtered to obtain a median of 5 million reads per sample. Using ENRICH software (*46*), read counts for each variant before and after selection were used to determine mating efficiency and invasion ability of *STE12* variants. Briefly, counts for a particular variant in the input and output libraries were normalized by their respective read totals to determine frequency in each, and a ratio of the output and input frequencies determine a variant’s functional score. Finally, enrichment scores are normalized by the enrichment of wild-type Ste12 in each selection. Treatment scores are calculated identically, and in all cases where difference scores are shown, the score represents log2(treated) – log2(untreated).

### Calculating intramolecular epistasis scores

Intramolecular epistasis scores were defined as the deviation of double-mutant’s functional score (*W*_*ij*_) from the multiplied scores of its constituent single-mutants (*w*_*i*^*^_ *w*_*j*_). A negative epistasis score indicates that the deleterious effect of one mutation is increased by the presence of the partner mutation, while a positive epistasis score indicates that the partner mutation decreases the deleterious effects of individual mutations(*47*).

### Quantitative mating assay

Individual variants tested for mating efficiency were treated identically to the large-scale mating selection experiments, except genotypes were scored individually on selective plates for either mated diploids (2N) or both diploids and unmated MATa haploids (2N, 1N). The proportion of mated individuals (mating efficiency) is taken as the ratio of colony counts on diploid to counts on diploid + haploid plates (2N/2N1N). Ste12 variants were always tested alongside wild-type to determine relative changes in response to treatment.

### STE12 variant RNA-seq

RNA was extracted from yeast cells harboring the *STE12* mutant library grown under non selective conditions using acid phenol extraction as previously described(*48*). *STE12*-specific cDNA was created using a gene-specific cDNA primer and Superscript III (Life Technologies). cDNA was amplified in a manner similar to the plasmids, and prepared for sequencing using Illumina Nextseq.

### Large-scale analysis of Ste12 binding sites *in vivo*

A *HIS3* reporter gene was used for testing large populations of binding site variants (Fig. 5 - Suppl. Fig. 2) in media lacking histidine. Although Ste12 does bind single PREs as a monomer, two sites are needed for signal detection in reporter assays (*49*). We designed oligonucleotides that maintain the central portion of the native PRE, but randomized six surrounding bases on either side (NNNNNNTTTCAAAATGAAANNNNNN). This library was cloned into a promoter from the same plasmid used in the luciferase assay, at the same position as the native PRE. Strains containing one of four Ste12 protein variants were transformed with the same binding site library reporter population, and grown overnight in synthetic media lacking histidine with 10mM of the His3 competitive inhibitor 3-amino-triazole. Three biological replicate selections were conducted for each Ste12 protein variant tested against the binding site library. Cells were collected and sequenced before and after selections to determine enrichment scores for each binding site variant. Computational analysis of binding site enrichment scores was identical to pipeline for Ste12 protein variants. Enrichment of all binding site variants are shown relative to the enrichments of empty plasmids, which were spiked-in to the binding site library as control. The 250 binding sites with the highest enrichments were grouped and Weblogo (*50*) was used to generate base preference plots.

### Evolutionary analysis of Ste12 DNA-binding domain among fungi

Using fungal genomes deposited into NCBI and from the fungal sequencing project at Joint Genome Institute, we created a BLAST database of translated coding sequences and queried with the *S. cerevisiae* Ste12 protein sequence. Full protein sequences from BLAST hits were aligned using MUSCLE (*51*) and used to determine conservation at each site in the mutated segment. The secondary structure of *S. cerevisiae*’s Ste12 DNA-binding domain was determined using Psipred (*52*).

### Analysis of Ste12 ChIP data

Genes bound by Ste12 only in mating conditions were compared to those bound by Ste12 only in invasion conditions (*14*). FIMO(*53*) was used to extract all matches to the Ste12 binding site in each of these gene sets. The frequency of the each possible base at position 6 of the core Ste12 motif was determined relative to the genomic background (Ste12 binding sites in all upstream sequences, no separation by mating or invasion gene function). The core 6 base frequencies relative to genomic background was then compared for genes bound in mating or invasion conditions. We used the same method to look at core 6 base frequencies at genes bound by both Ste12 and Tec1 or those bound by Ste12 alone during invasion.

## Acknowledgments

We thank J. Thomas for advice on analysis of fungal variation. We thank D. Fowler and H. Malik for comments on the manuscript. The work was supported by NIH giant R01GM114166 to C.Q. and S.F. M.W.D. was supported by an NSF Graduate Research Fellowship and a WRF-Hall fellowship. S.F. is an investigator of the Howard Hughes Medical Institute, which supported J.T.C.

## Author Contributions

M.W.D., J.T.C. S.F., and C.Q. conceived and interpreted experiments. M.W.D., S.F., and C.Q. wrote the manuscript. J.T.C. and J.A.C. provided comments. M.W.D. conducted experiments and data analysis: J.A.C. contributed to high-throughput binding assay.

## Data Availability

High-throughput sequencing reads have been submitted to NCBI SRA. awaiting accession number. Datasets with calculated enrichment scores for each tested variant in each condition, along with positional mean scores, and scores from binding site library selections are provided in source data files.

---

**Fig. 1 - Suppl. Fig 1.**
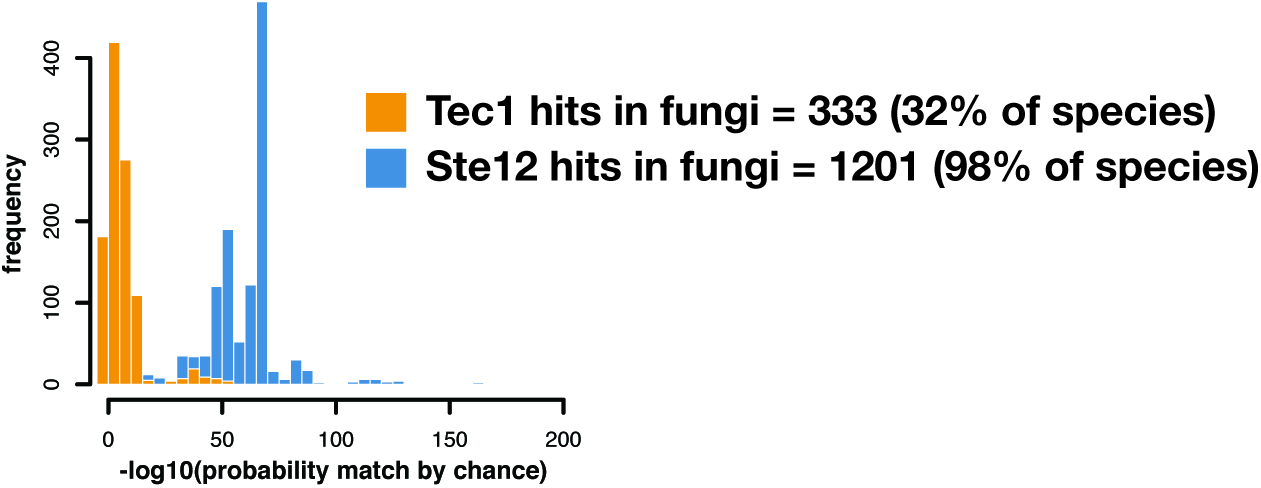
*TEC1* is frequently absent in fungi whereas *STE12* is maintained. We queried a protein database of 1229 fungal genomes and identified matches to either *S. cerevisiae* Tec1 (orange) or Ste12 (blue) sequence.

**Fig. 2 - Suppl. Fig 1.**
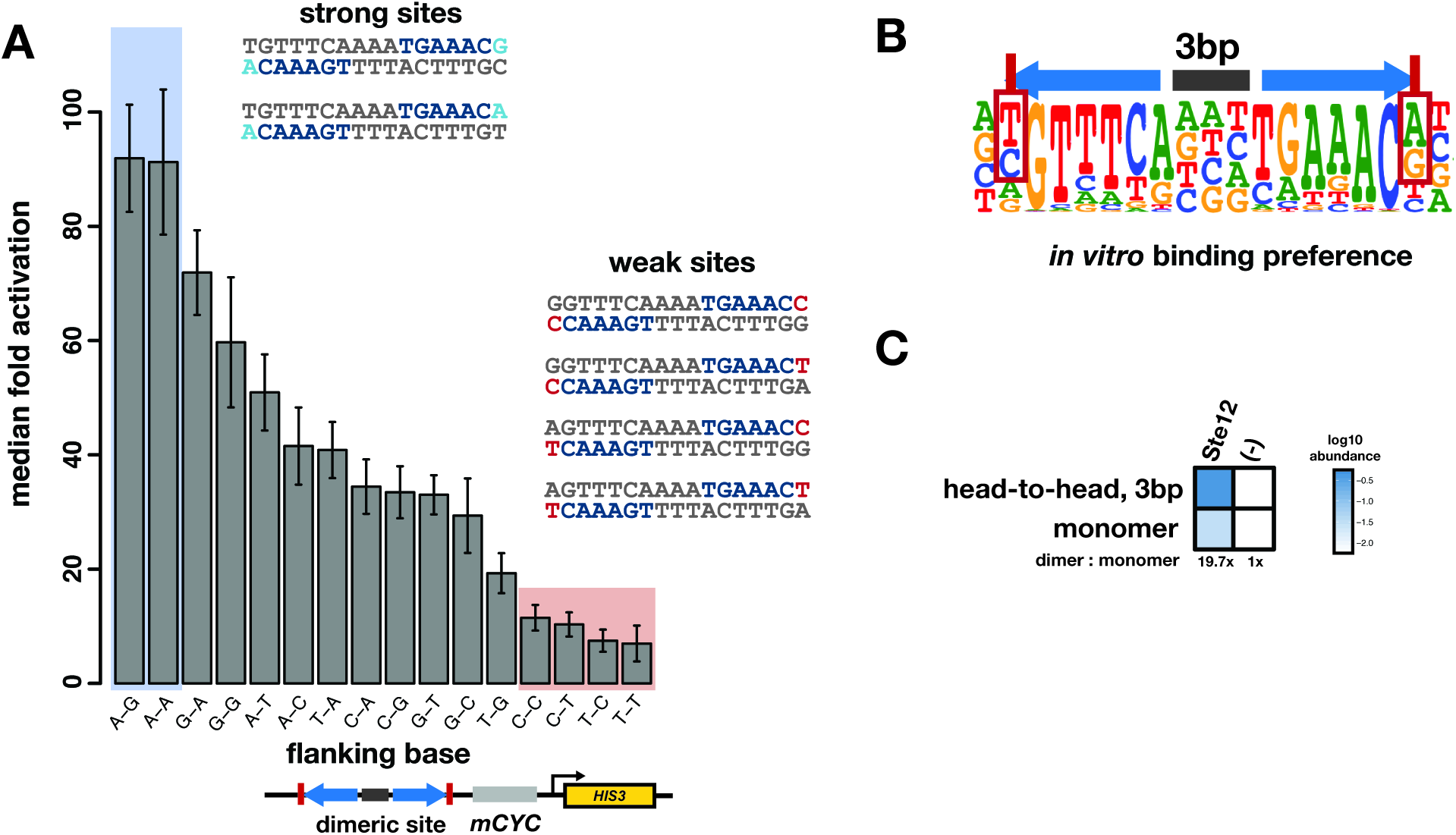
Wild-type Ste12’s flanking base preferences *in vitro* are correlated with basal activity *in vivo*. (A) We collected activity scores among all sites containing perfect dimer sites GTTTCANNNTGAAAC, separated by the combination of flanking bases following the final base of the core motif (n > 50 for all flank combinations), and plotted the median activation of these sets of sites. Nearly an order of magnitude of Ste12’s activation can be explained by the combination of flanking bases on a dimeric site. (B) Ste12’s flanking base preferences identified *in vitro* by HT-SELEX (from Fig. 2B). (C) Ste12 was unable to activate a monomeric site *in vivo*, and this preference is reflected in its *in vitro* preference for dimeric binding.

**Fig. 2 - Suppl. Fig 2.**
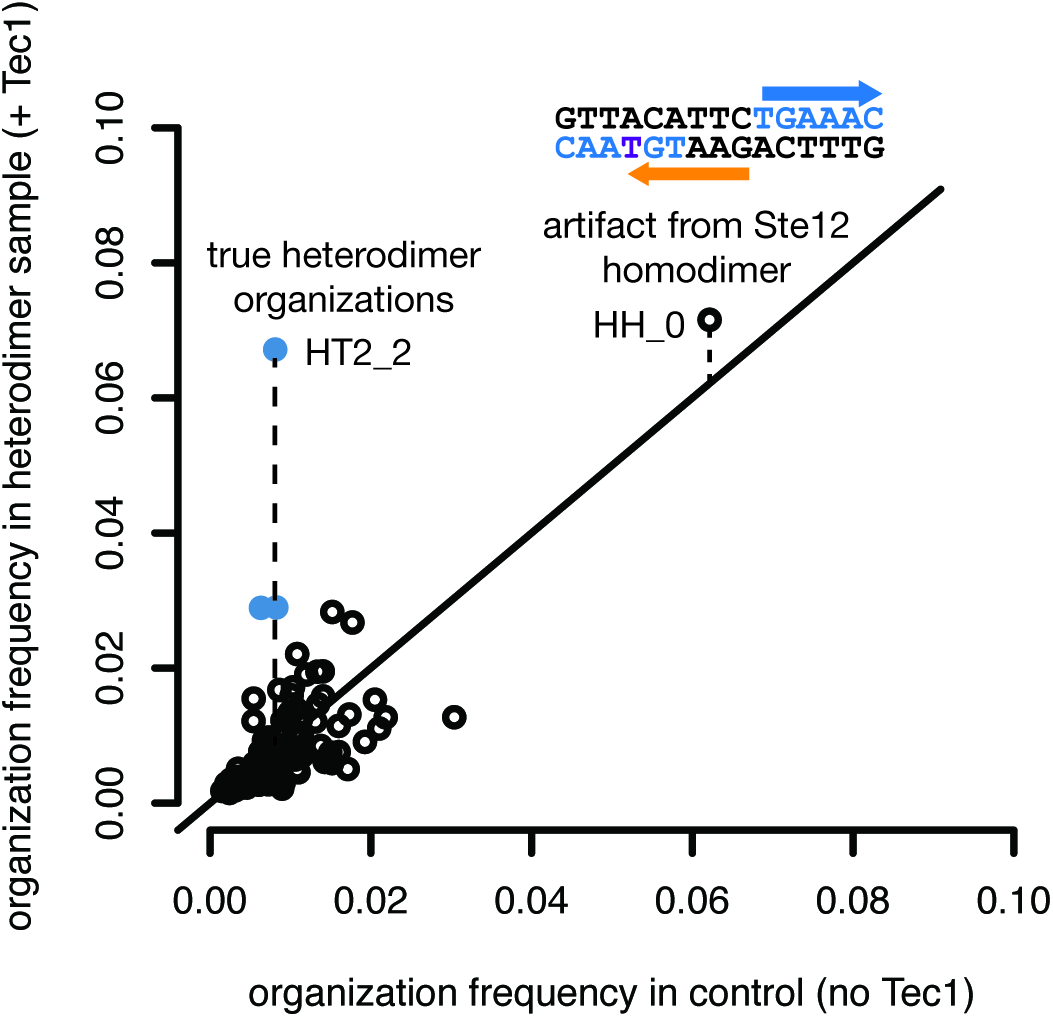
Comparing co-binding sample with Ste12 alone sample allows removal of false Tec1 sites. Each point represents a particular spacing and orientation combination. The frequency of each combination found in output SELEX samples is shown for Ste12 alone (x-axis) and for Ste12 + Tec1 (y-axis). The most frequent organization in the Ste12 + Tec1 sample is head-to-head 0bp, but this site arises from fortuitous spacer sequences between Ste12 homodimer sites that resemble a Tec1 site (example indicated above head-to-head 0bp point). Normalizing each organization by its representation in the Ste12 alone sample reveals the true heterodimeric sites (colored in blue). (HH_0 = head-to-head, 0bp spacer, HT2_2 = head-to-tail Tec1 before Ste12, 2bp spacer)

**Fig. 3 - Suppl. Fig 1.**
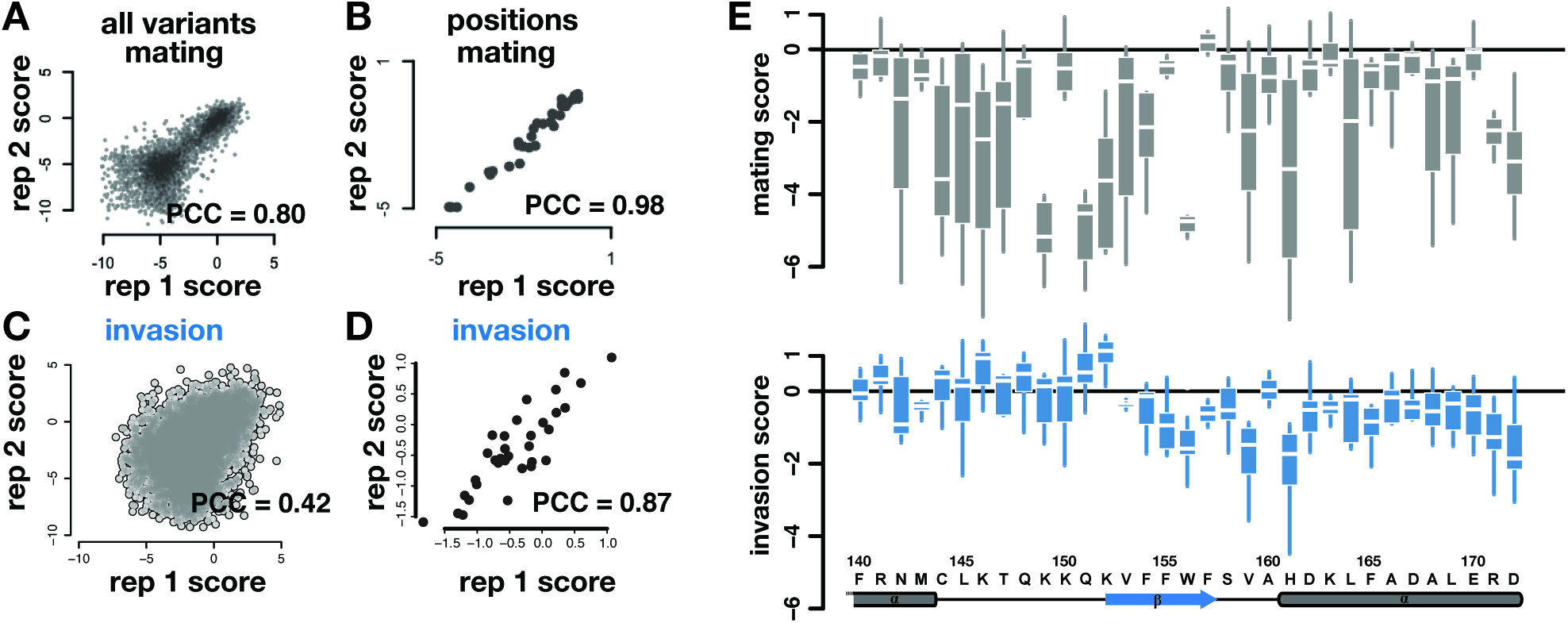
Mating and invasion enrichment scores are reproducible across replicate experiments. Selection for each trait was done in three biological replicates. We computed Pearson correlation coefficients (PCC) between replicates for all variants (A,C) as well as positional means (B,D). Shown are correlations for replicate 1 and 2, results were similar for other replicate pairs. Invasion selections were less well correlated. Nevertheless, positional mean scores were strongly correlated for both traits across replicates. (E) Variability in per-position effect of mutation is shown as boxplots of all mutations tested at each site for mating (grey) and invasion (blue).

**Fig. 3 - Suppl. Fig 2.**
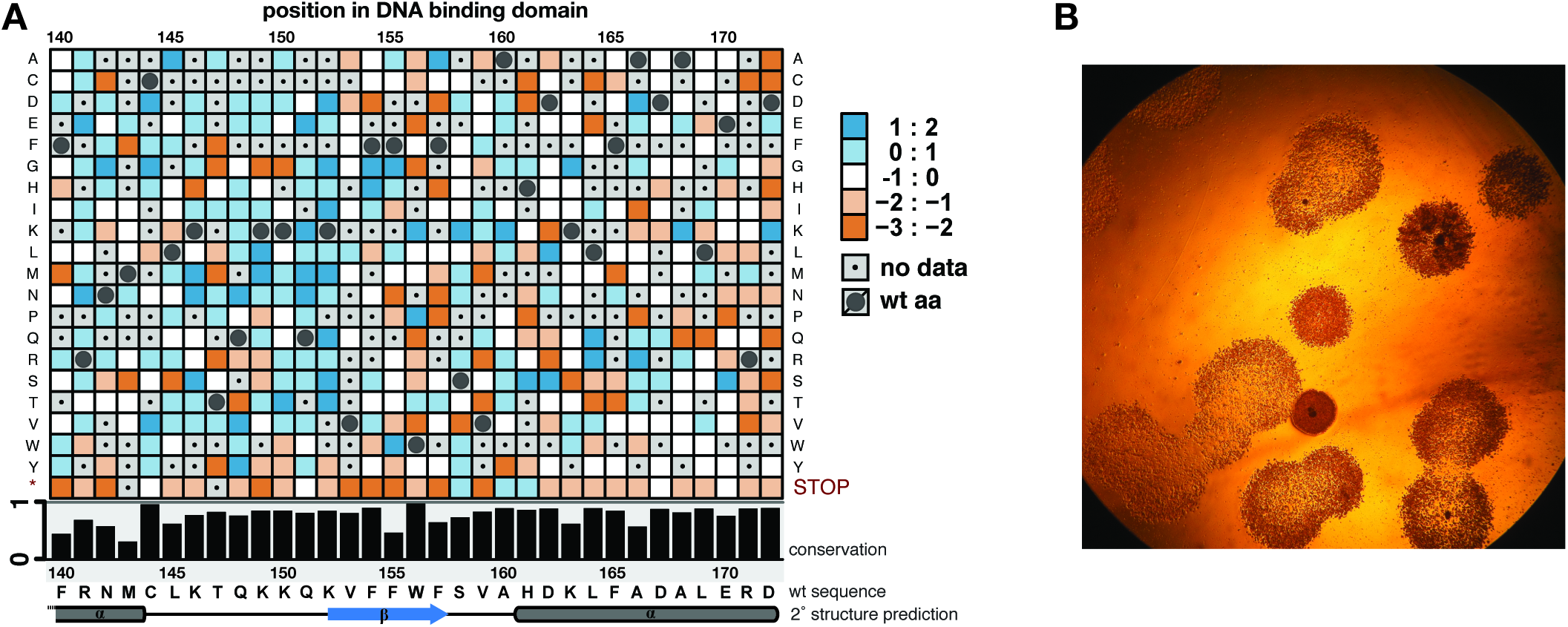
Variation in the highly conserved DNA binding domain of Ste12 generates a range of invasion phenotypes. (A) A heatmap of invasion scores for all single mutants is displayed as in Fig. 3B for mating scores. (B) A 10x magnified image of a washed plate containing invaded colonies with different Ste12 variants shows phenotypic variation in invasion.

**Fig. 3 - Suppl. Fig 3.**
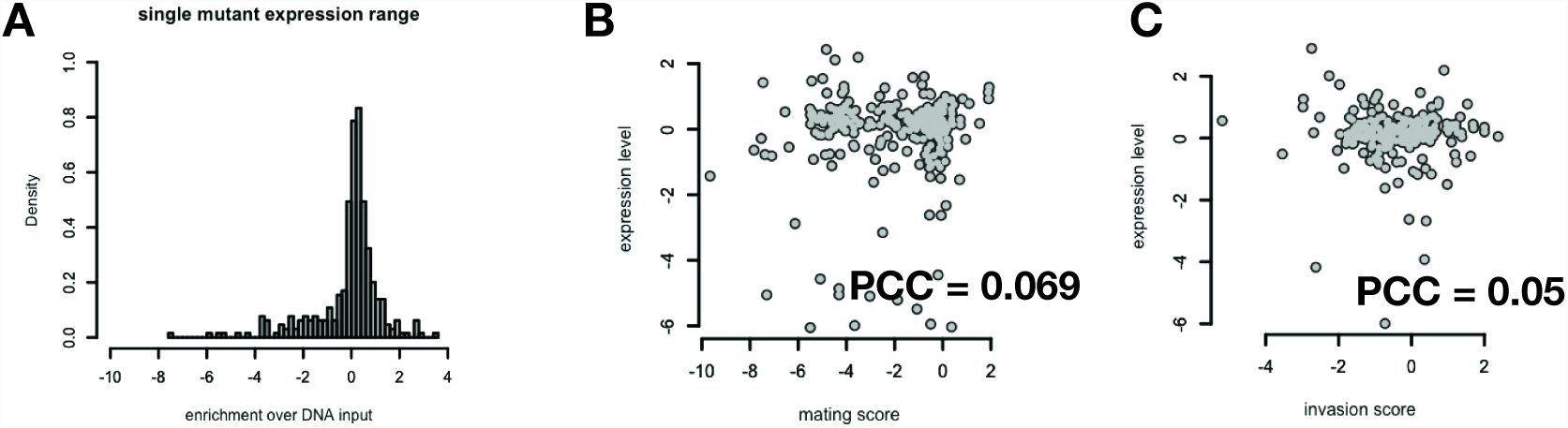
Variant expression shows little variation and no correlation with phenotypic effects. (A) The distribution of log2 enrichment of transcripts relative to plasmid counts for each Ste12 variant rarely deviates from zero, indicating low variation in expression level among variants. Each variant’s expression level is plotted against each variant’s trait enrichment score, for (B) mating and (C) invasion, demonstrating that variant expression levels did not affect phenotype.

**Fig. 4 - Suppl. Fig 1.**
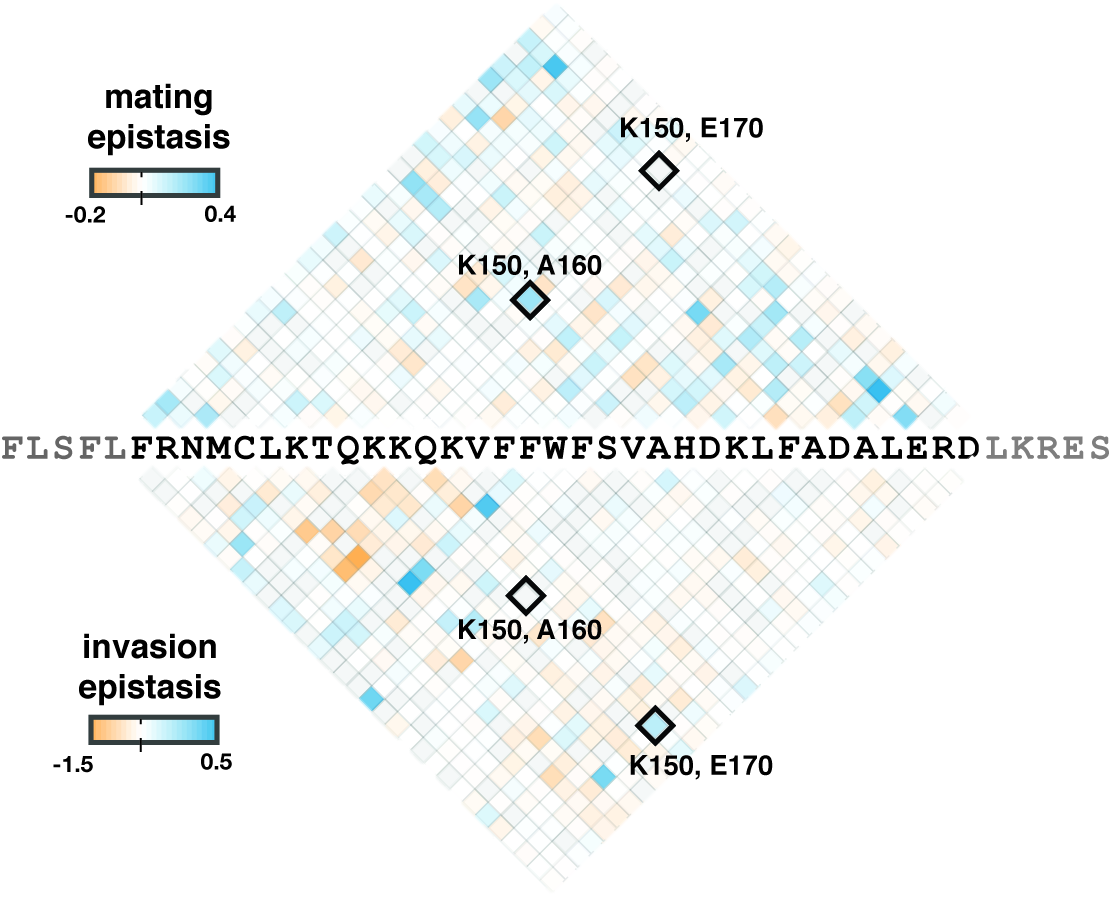
Double mutant analysis identifies pairs of positions with strong epistasis, and such pairs differ between mating and invasion. For each double mutant, we calculated epistasis as the deviation of that mutant’s effect from the multiplied effect of its constituent single mutants (see Methods). Scores are displayed as in Fig. 2A; epistasis scores across all combinations of mutations tested at each pair of sites was used to calculate the displayed mean epistasis score. Sites with strong positive (shades of blue) or negative epitasis scores (shades of orange) differ between traits. Strong intramolecular epistasis occurs more frequently between residues in close proximity, suggesting that Ste12 conformation differs between both traits. Two pairs with trait-specific epistatic interactions are indicated in boxes.

**Fig. 4 - Suppl. Fig 2.**
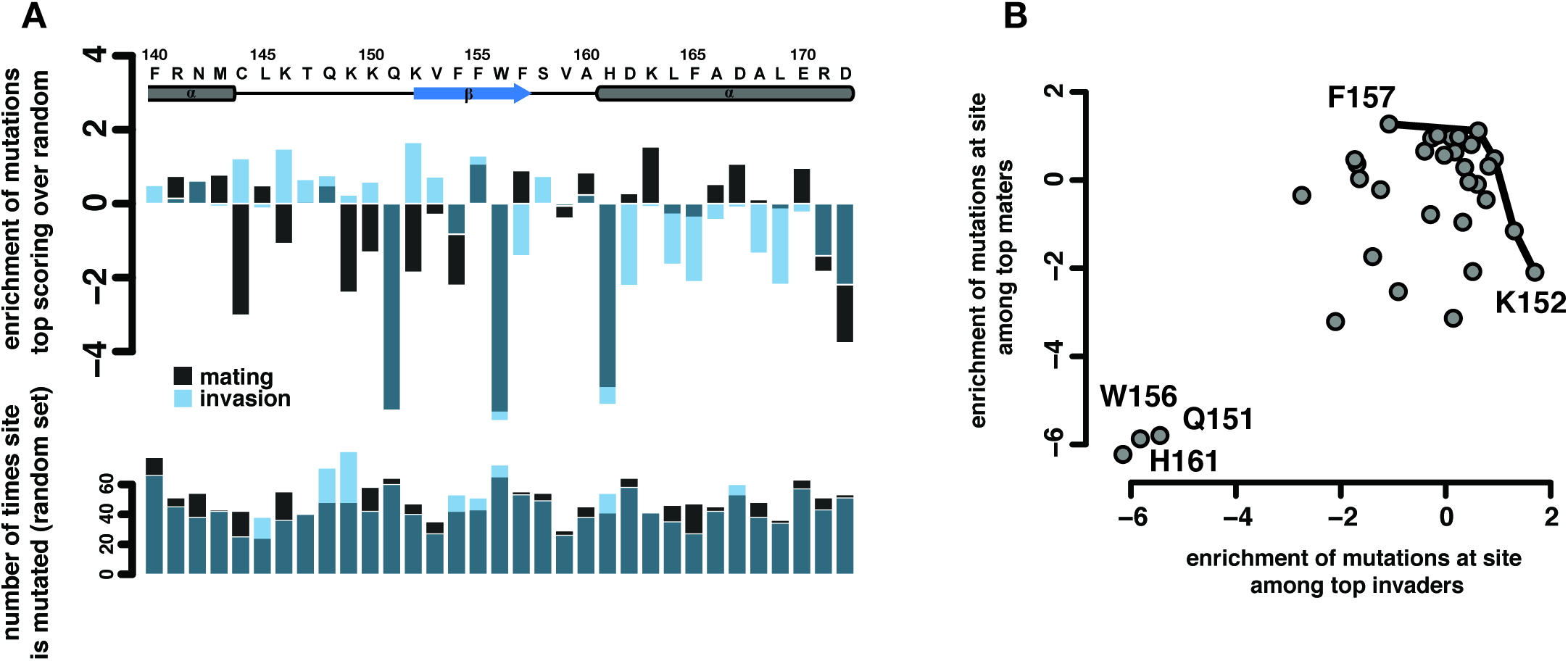
Orthogonal analysis reveals positions contributing to ‘separation-of-function’ between mating and invasion. >Per-site mutation frequencies were calculated for the top 750 mating variants, top 750 invasion variants, and a random set of 750 variants from each dataset. (A) Enrichment of per-site mutations for top performing variants in each trait recapitulates bipartite arrangement of effects (‘separa-tion-of-function’), as well as sites increasing one trait at the cost of the other (e.g. see K152 and F157, tradeoff indicated with blue/black bars). (B) Per-site enrichment scores identify similar sites on the empirical Pareto front (Fig. 4B), as well as the same sites in which mutations are deleterious in both traits.

**Fig. 4 - Suppl. Fig 3.**
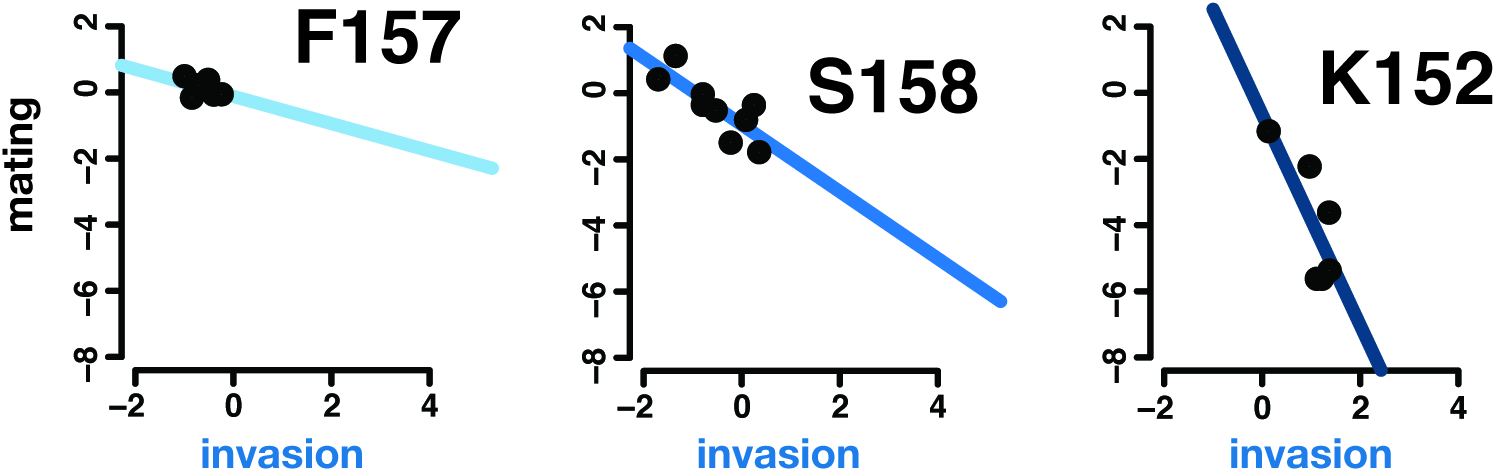
Individual mutations tested at sites critical for separation of function between mating and invasion. Mating and invasion scores for all single amino acid change mutations at each indicated position were plotted, and a simple linear model was fitted. Steeper slopes indicate that mutations at that position are more likely to shift trait preference toward invasion.

**Fig. 4 - Suppl. Fig 4.**
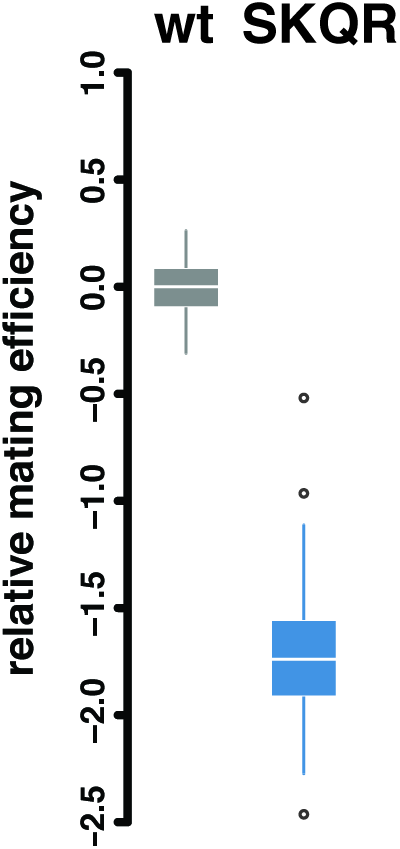
SKQR mutant shows decreased mating efficiency. Because the SKQR variant was not present in our initial Ste12 variant library, we used a standard quantitative mating assay to determine its mating efficiency relative to wild-type Ste12 as in Fig. 4F.

**Fig. 4 - Suppl. Fig 5.**
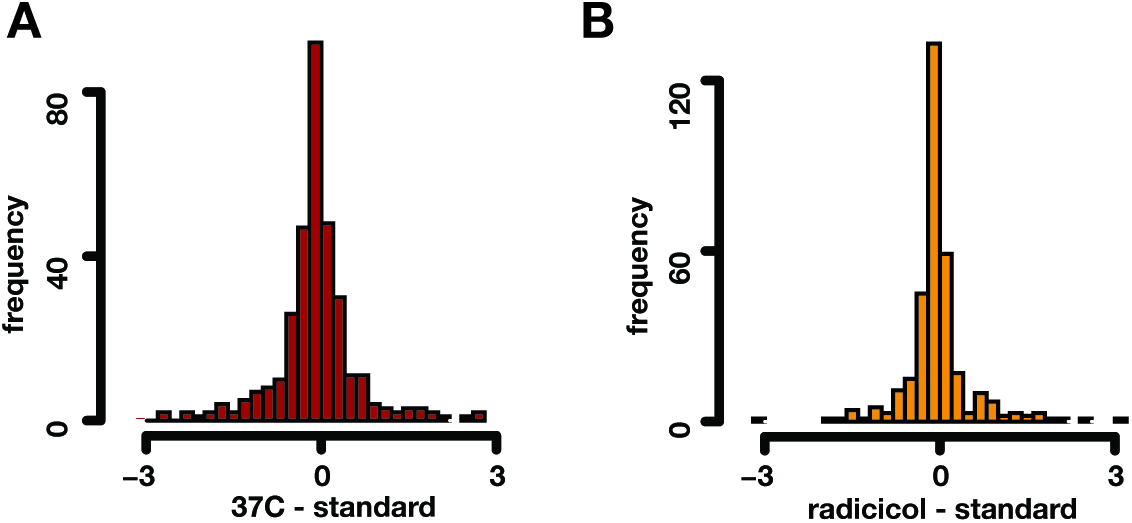
The majority of Ste12 variants respond like wild-type Ste12 to increased temperature or Hsp90 inhibition. The distribution of differences of log2 mating scores for all single mutants (treated - untreated) are plotted for high-temperature (red, A) and Hsp90-reduced (orange, B) conditions.

**Fig. 5 - Suppl. Fig 1.**
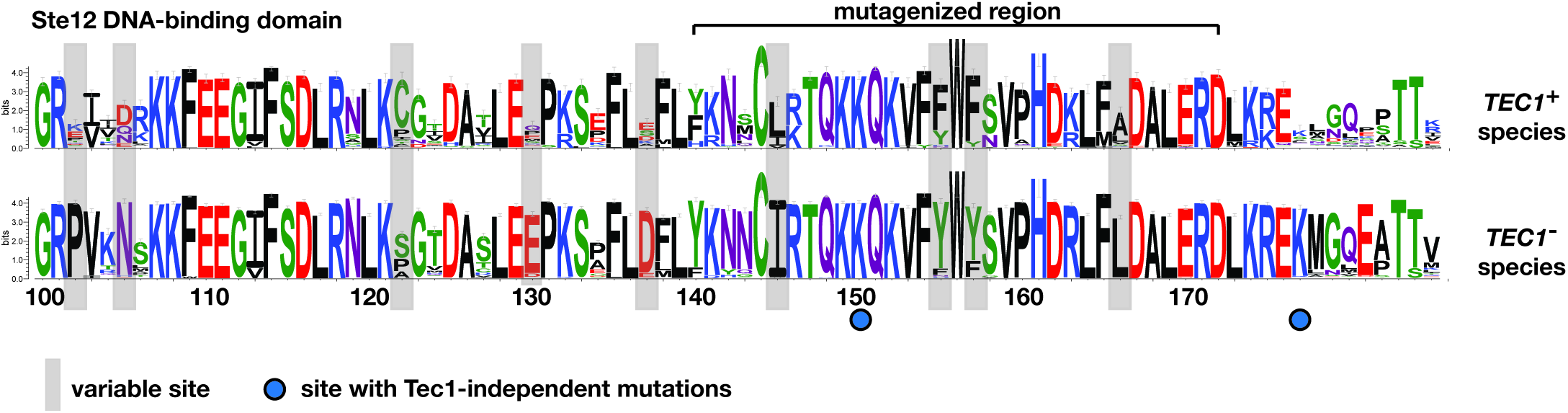
Variation in Ste12’s DNA-binding domain is associated with presence of *TEC1* gene. A weblogo of Ste12’s DNA-binding domain was generated as in Figure 1D, except that the fungal species were split into two groups: those whose genome contains a copy of *TEC1* (n=333) and those whose do not (n=715).

**Fig. 5 - Suppl. Fig 2.**
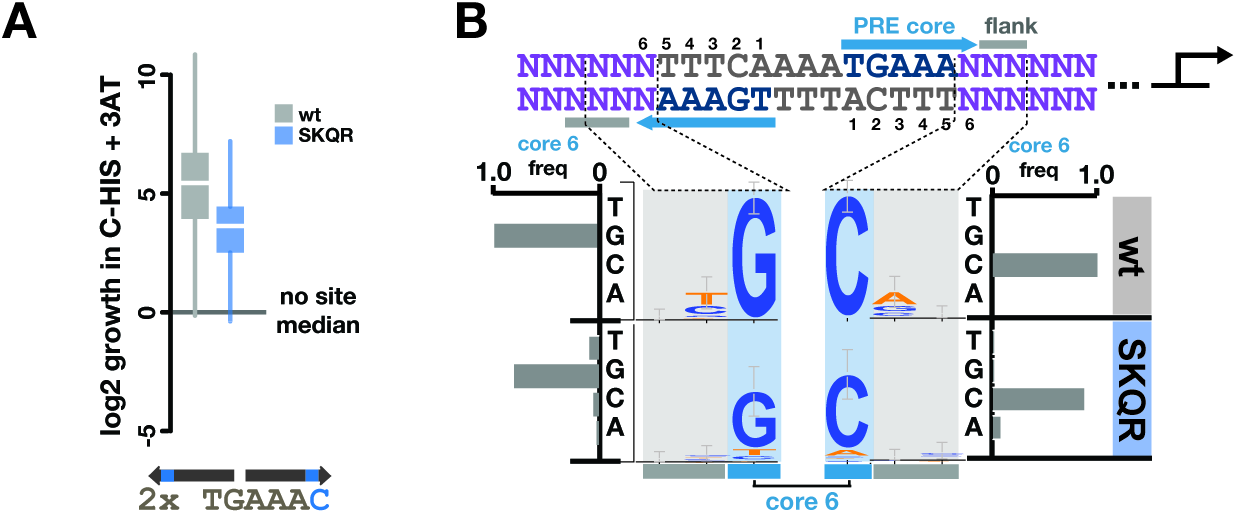
SKQR shows reduced activation at perfect match dimeric sites compared to wild-type Ste12, but can activate degenerate sites. A library of binding sites was cloned into a HIS3 reporter gene and used in growth selections in the BY4741 *Ste12Δ* background. (A) The activity of wild-type Ste12 and SKQR variant with all binding sites containing two canonical PREs is shown as a boxplot (n = 725, 747). (B) The *in vivo* sequence preferences for wild-type Ste12 and SKQR are shown as logo plots of three positions (core position 6 and two flanking bases, no preferences were seen at further flanking bases) of each PRE. Grey error bars on logo plots represent 95% confidence interval at each base. Bar charts show the frequency of base preference at core position 6.

